# Neuroanatomy Reflects Individual Variability in Impulsivity in Youth

**DOI:** 10.1101/2025.04.23.650222

**Authors:** Elvisha Dhamala, Erynn Christensen, Jamie L. Hanson, Jocelyn A. Ricard, Noelle Arcaro, Simran Bhola, Lisa Wiersch, Katharina Brosch, B.T. Thomas Yeo, Avram J. Holmes, Sarah W. Yip

**Author notes:** Corresponding author: Elvisha Dhamala.

## Abstract

Individual differences in neural circuits underlying emotional regulation, motivation, and decision-making are implicated in many psychiatric illnesses. Interindividual variability in these circuits may manifest, at least in part, as individual differences in impulsivity at both normative and clinically significant levels. Impulsivity reflects a tendency towards rapid, unplanned reactions to internal or external stimuli without considering potential negative consequences coupled with difficulty inhibiting responses.

Here, we use multivariate brain-based predictive models to explore the neural bases of impulsivity across multiple behavioral scales, neuroanatomical features (cortical thickness, surface area, and gray matter volume), and sexes (females and males) in a large sample of youth from the Adolescent Brain Cognitive Development (ABCD) Study at baseline (n=9,099) and two-year follow-up (n=6,432).

Impulsivity is significantly associated with neuroanatomical variability, and these associations vary across behavioral scales and neuroanatomical features. Impulsivity broadly maps onto cortical thickness in dispersed regions (e.g., inferior frontal, lateral occipital, superior frontal, entorhinal), as well as surface area and gray matter volume in specific medial (e.g., parahippocampal, cingulate) and polar (e.g., frontal and temporal) territories. Importantly, while many relationships are stable across sexes and time points, others are sex-specific and dynamic.

These results highlight the complexity of the relationships between neuroanatomy and impulsivity across scales, features, sexes, and time points in youth. These findings suggest that neuroanatomy, in combination with other biological and environmental factors, reflects a key driver of individual differences in impulsivity in youth. As such, neuroanatomical markers may help identify youth at increased risk for developing impulsivity-related illnesses. Furthermore, this work emphasizes the importance of adopting a multidimensional and sex-specific approach in neuroimaging and behavioral research.

## Introduction

Impairments in emotional regulation, motivation, and decision-making are prevalent across a range of psychiatric illnesses^1^ and often emerge during early adolescence^2,3^. These impairments contribute to the heterogeneity observed within psychiatric illnesses and may initially appear as more fundamental alterations in processes and behaviors such as impulsivity^4,5^. Impulsivity is a multifaceted construct that reflects “a predisposition toward rapid, unplanned reactions to internal or external stimuli without regard to the negative consequences of these reactions to the impulsive individual or to others.^6^” Importantly, changes in impulsivity are a normal part of development. However, in some individuals, heightened levels of impulsivity may indicate increased risk for psychiatric illness^7–9^ (see Supplemental Materials for specific examples).

Although often treated as a single construct, the term impulsivity encompasses a variety of distinct but related functions that promote impulsive behavior^10^. These include an individual’s (in)ability to consider the consequences of a behavior (lack of premeditation), tendency to disengage from tasks due to boredom or difficulty before completion (lack of perseverance), responses to emotional states (positive and negative urgency), and motivation to experience rewarding sensations (sensation seeking)^9^. Impulsivity can also be considered a product of two systems that promote impaired self-regulation: the behavioral inhibition and approach systems (BIS/BAS)^11^. The BIS prevents actions that may lead to a negative outcome^12^, while the BAS encapsulates sensitivity to, and motivation for, reward/punishment, as well as escape from punishment, therefore encouraging incentive-motivated behavior^12^. These conceptualizations highlight the variability in how impulsivity is defined and measured, posing a challenge for research, and underscoring the need for greater conceptual clarity across different definitions of impusivity^6,10^. Consequently, a crucial consideration in exploring the neuroanatomical basis of impulsivity is the extent to which different, yet related, components of this construct may be subserved by shared versus distinct neurobiological substrates.

Corticolimbic and corticostriatal circuitry regulate impulsivity^8,13–16^. The corticolimbic system contributes to processing emotional salience and regulating emotional responses^17^, while the corticostriatal system is involved in motivated behavior, reward processing, learning, and habit formation^18^. These systems mature throughout development^19^ and this is accompanied by significant changes in synaptic connectivity and myelination^20–22^. Critically, this development is asynchronous. Limbic and striatal structures mature earlier than cortical structures, including the prefrontal cortex, resulting in heightened impulsivity during this period of developmental ‘mismatch’^21,23,24^. Behaviorally, emotional regulation, motivation, and impulse control evolve throughout development with rapid changes in early life followed by gradual changes during adolescence^25–29^. This change is paralleled by changes in their neural substrates beyond corticolimbic and corticostriatal circuits, such as the insula and cingulate cortex^7,29,30^. However, previous studies examining these relationships have often focused on establishing univariate, cross-sectional associations between specific brain circuits and individual behavioral measures in small sample sizes. In contrast, brain-based predictive models use machine learning to analyze whole-brain multivariate relationships between brain features and behavioral measures^31^, which account for the interconnected nature of the brain, unlike traditional analyses focused on single regions. These models along with the large sample sizes provided by big data initiatives can provide insights into the whole-brain neuroanatomical basis of impulsivity^32^. In addition, longitudinal data can be used to examine whether these relationships are consistent throughout development. Here, we leverage these approaches to garner insights into the neural substrates of impulsivity in youth.

Importantly, neurodevelopmental processes and behavioral expressions vary between males and females, raising questions about the extent to which sex-specific neuroanatomical patterns contribute to observed differences in impulsivity. There are significant sex differences in the developmental trajectories of corticolimbic and corticostriatal systems^33,34^, although findings have not always been consistent across studies. As an example, on average, females have greater relative volume in the prefrontal and orbitofrontal regions, while males have greater volume in ventral temporal and occipital regions^35^. Similarly, sex differences have been reported in impulsivity, but these results are also inconsistent^36^. Thus, it is plausible that sex differences exist across neuroanatomy, impulsivity, and their interrelationships, highlighting the importance of considering sex differences when studying impulsivity, particularly within a developmental framework. Furthermore, within the context of recently established large data initiatives, it remains to be determined whether sex differences in impulsivity are driven by unique neuroanatomical substrates. This can be addressed by establishing sex-specific relationships between neuroanatomy and impulsivity and examining the extent to which they overlap.

Cortical thickness (CT), surface area (SA), and gray matter volume (GMV) reflect different aspects of neuroanatomy. CT (i.e., distance between the brain’s outer surface and gray-white matter junction) reflects neuronal density and arrangement^37,38^. It increases rapidly during the prenatal period, continues growing after birth, peaks in early childhood (with regional variation), and then gradually thins^39^. Cortical SA (i.e., area of the pial surface) is linked to the organization and complexity of cortical columns ^37,38^ as well as neuronal proliferation^37,38^ and gyrification^40^. SA expands prenatally and through childhood, peaking in late childhood/early adolescence, and then gradually declines^39^. GMV, encompassing thickness and SA, reflects the total amount of cells and synapses^37,38^, and generally follows the same developmental trajectory observed for SA^39^. Changes in these neuroanatomical features result from neurogenesis, synaptogenesis, synaptic pruning, cell death, and alterations in cell size and density, and are linked to various psychiatric conditions^37,38^. Given these complexities, a multimodal analysis considering all three features of brain structure across all regions of the brain is warranter to reveal their unique and shared contributions to impulsivity, facilitating a more holistic understanding of these relationships.

Here, we investigated the sex-specific neuroanatomical basis of impulsivity, across different neuroanatomical features and impulsivity measures, in a large sample of youth from the Adolescent Brain Cognitive Development (ABCD) Study at baseline and two-year follow-up. We show that neuroanatomical features are associated with impulsivity and there are notable sex differences in these relationships. We also demonstrate that different domains of impulsivity are linked to shared and distinct neuroanatomical features. Some features vary across facets of impulsivity, others across sexes, and others across time points. These findings highlight substantial individual variability in the neural basis of impulsivity in youth. Understanding these distinct markers of impulsivity is crucial for establishing normative developmental patterns and paves the way for development of more effective early interventions grounded in neurobiological mechanism to prevent psychiatric illness.

## Methods

An overview of the methods is provided below. Details are in the Supplemental Materials.

### Dataset

The ABCD Study is following a large community-based sample of children and adolescents throughout the course of development^41^. Participants are assessed on a comprehensive set of neuroimaging, behavioral, developmental, and psychiatric batteries. In this study, we used imaging and impulsivity data from 9,099 participants at baseline (ages 9-10 years) and 6,432 participants at the two-year follow-up (see **Table S1** for demographic data). Details on data inclusion procedures (e.g., exclusion criteria for neuroimaging data) are provided in the Supplemental Materials (see **Figures S1-S2** for our participant inclusion pipeline).

### Neuroimaging

The neuroimaging protocol and specific parameters for T1-weighted scans are detailed in previous publications^41,42^. We used measures of CT (mean), SA (total), and GMV (total) for 68 cortical regions (34 per hemisphere) from the Desikan-Killiany parcellation as provided on the NIMH Data Archive. Regional SA and GMV measures, but not CT, were proportionally corrected for individual differences in intracranial volume by dividing the raw values by intracranial volume, as recommended by prior work^43^ (see **Figure S3** for average measures and **Figure S4** for sex differences in the measures).

### Impulsivity

Impulsivity-related measures were derived from the Behavioral Inhibition/ Activation System (BIS/BAS) and the Modified Urgency, (lack of) Planning (or Premeditation), (lack of) Perseverance, Sensation-Seeking, and Positive Urgency (UPPS-P^44^) Short Version scales.

### Differences in Impulsivity Across Development and Across Sexes

We used non-parametric Mann-Whitney U rank tests to evaluate differences in the behavioral measures (4 BIS/BAS, 5 UPPS-P) between baseline and two-year follow-up as the scales were not normally distributed. We also used non-parametric Mann-Whitney U rank tests to evaluate sex differences in the behavioral measures at each time point. We corrected p-values for multiple comparisons within each behavioral scale using the Benjamini-Hochberg False Discovery Rate (q=0.05) procedure^45^. We also computed sex-independent and sex-specific full correlations between the measures at each time point to evaluate their co-expression in youth.

### Predictive Modeling

We used a cross-validated brain-based predictive modeling framework^31^ which we have leveraged in prior work^43,46–50^. This framework avoids data leakage and minimizes overfitting to capture robust, reliable, and interpretable associations between imaging-derived measures and phenotypic data. For each pair of neuroanatomical features and behavioral measure, we developed separate sets of sex-independent (i.e., including the entire sample) and sex-specific (i.e., in either females or males) linear ridge regression models at each of the two time points to predict impulsivity based on neuroanatomy. We trained each model on neuroanatomical features (either CT, SA, or GMV) from 68 regions to predict a single impulsivity measure. We quantified model performance using prediction accuracy^50–53^ and assessed significance in comparison to null distributions. We corrected p-values for multiple comparisons within each behavioral scale using the Benjamini-Hochberg procedure^45^.

### Feature Weights

For models yielding reliable brain-behavior relationships in both sexes (as compared to null distributions), we conducted a series of analyses to determine feature importance and to guide mechanistic understanding. This conservative approach ensures that comparisons are only made where meaningful brain-behavior relationships are present in both sexes. We transformed the feature weights obtained from the models using the Haufe transformation^54^ (to increase their interpretability and reliability^52,55,56^) and then calculated a mean feature importance for each set of models. We computed cosine similarities between the mean feature importance values to evaluate overlap in the regional features associated with different impulsivity measures.

## Results

### Expressions of impulsivity vary across youth

Analyses examining the distributions of the impulsivity measures and the correlations between them at baseline and two-year follow-up are presented in the Supplemental Materials (see **Figure S5** for results across all participants and **S6** for sex-specific results). These distributions are consistent with prior work examining impulsivity in the ABCD Study^57,58^, indicating that our sample is representative of the cohort. Broadly, these analyses indicated modest within-scale correlations of measures suggesting that they capture partially overlapping aspects of behavior, and significant but weak between-scale correlations indicating that the UPPS and BIS/BAS, while potentially related, measure somewhat independent constructs^59^.

### Neuroanatomy predicts impulsivity

Brain-based predictive models were used to quantify associations between neuroanatomy and impulsivity (**Figure 1, Table S2**).

**Figure 1:**
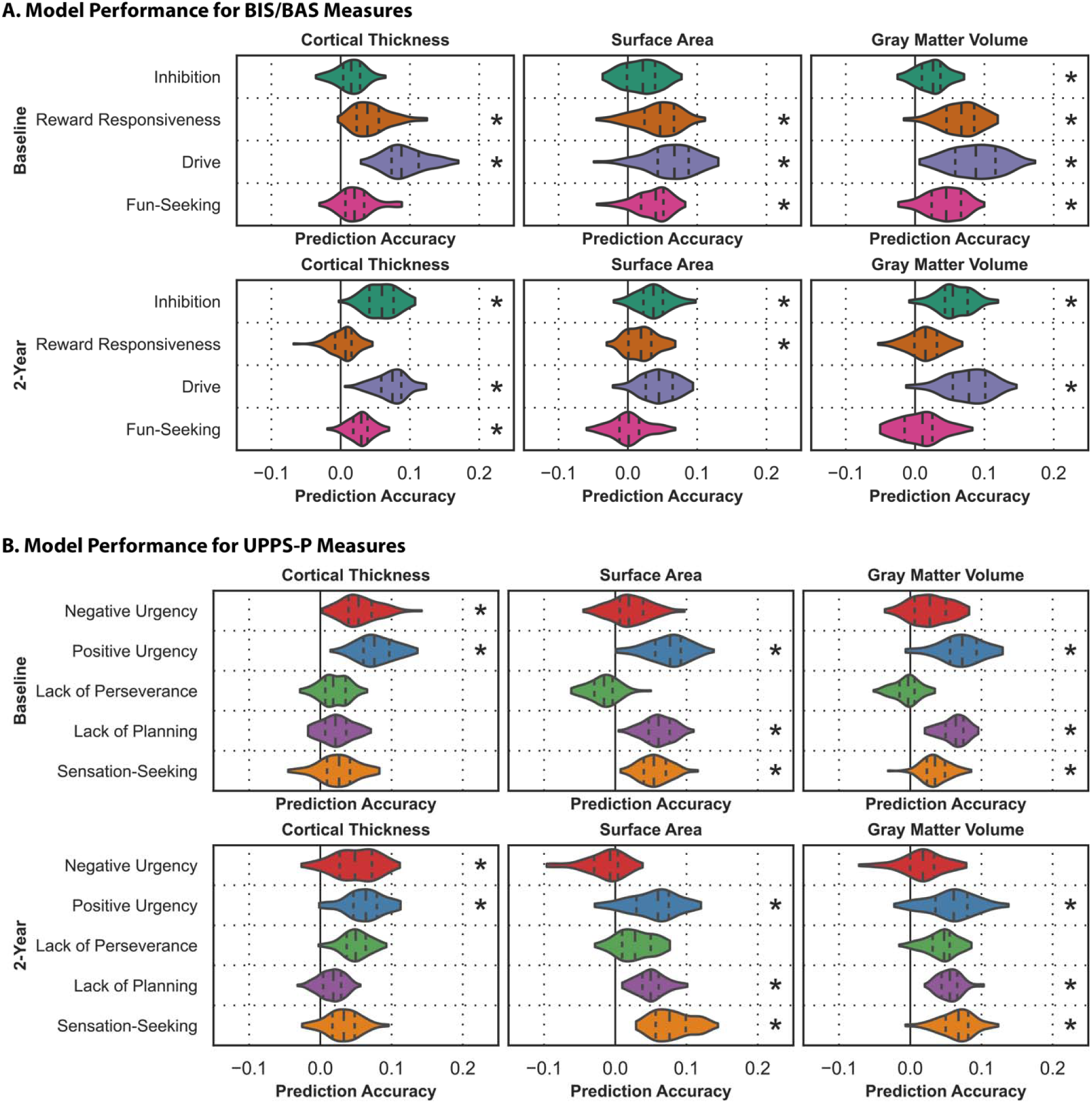
Neuroanatomy reflects individual differences in impulsivity. Prediction accuracies (correlation between observed and predicted values) for models trained to predict behavioral measures from the BIS/BAS (A) and UPPS-P (B) scales. Results for models based on CT (left), SA (center), and GMV (right) at baseline (top) and two-year follow-up (bottom) are shown. The shape of the violins indicates the distribution of values, the dashed lines indicate the median, and the dotted lines indicate the interquartile range. Asterisks indicate the model captured significant associations.

<u>CT:</u> Models based on CT accurately predicted reward-responsiveness (prediction accuracy, r=0.044, p_FDR_<0.001), drive (r=0.093, p_FDR_<0.001), negative urgency (r=0.055, p_FDR_=0.020), and positive urgency (r=0.078, p_FDR_=0.015) at baseline; and inhibition (r=0.059, p_FDR_=0.022), drive (r=0.073, p_FDR_=0.004), fun-seeking (r=0.029, p_FDR_=0.028), negative urgency (r=0.049, p_FDR_=0.008), and positive urgency (r=0.062, p_FDR_<0.001) at two-year follow-up.

<u>SA:</u> Models based on SA accurately predicted reward-responsiveness (r=0.043, p_FDR_<0.001), drive (r=0.064, p_FDR_=0.004), fun-seeking (r=0.035, p_FDR_=<0.001), positive urgency (r=0.074, p_FDR_<0.001), lack of planning (r=0.061, p_FDR_<0.001), and sensation-seeking (r=0.055, p_FDR_=0.003) at baseline; and inhibition (r=0.037, p_FDR_=0.044), reward-responsiveness (r=0.019, p_FDR_=0.044), positive urgency (r=0.055, p_FDR_=0.040), lack of planning (r=0.050, p_FDR_=0.040), and sensation-seeking (r=0.080, p_FDR_=0.025) at two-year follow-up.

<u>GMV:</u> Models based on GMV accurately predicted inhibition (r=0.024, p_FDR_=0.023), reward-responsiveness (r=0.065, p_FDR_=0.002), drive (r=0.088, p_FDR_=0.004), fun-seeking (r=0.043, p_FDR_=0.002), positive urgency (r=0.071 p_FDR_<0.001), lack of planning (r=0.062, p_FDR_<0.001), and sensation-seeking (r=0.036, p_FDR_=0.033) at baseline; and inhibition (r=0.058, p_FDR_=0.010), drive (r=0.077, p_FDR_<0.001), positive urgency (r=0.059, p_FDR_<0.001), lack of planning (r=0.056, p_FDR_=0.030), and sensation-seeking (r=0.065, p_FDR_=0.005) at two-year follow-up.

We next used the same framework to examine sex-specific associations (**Figure 2, Tables S3-S4**).

**Figure 2:**
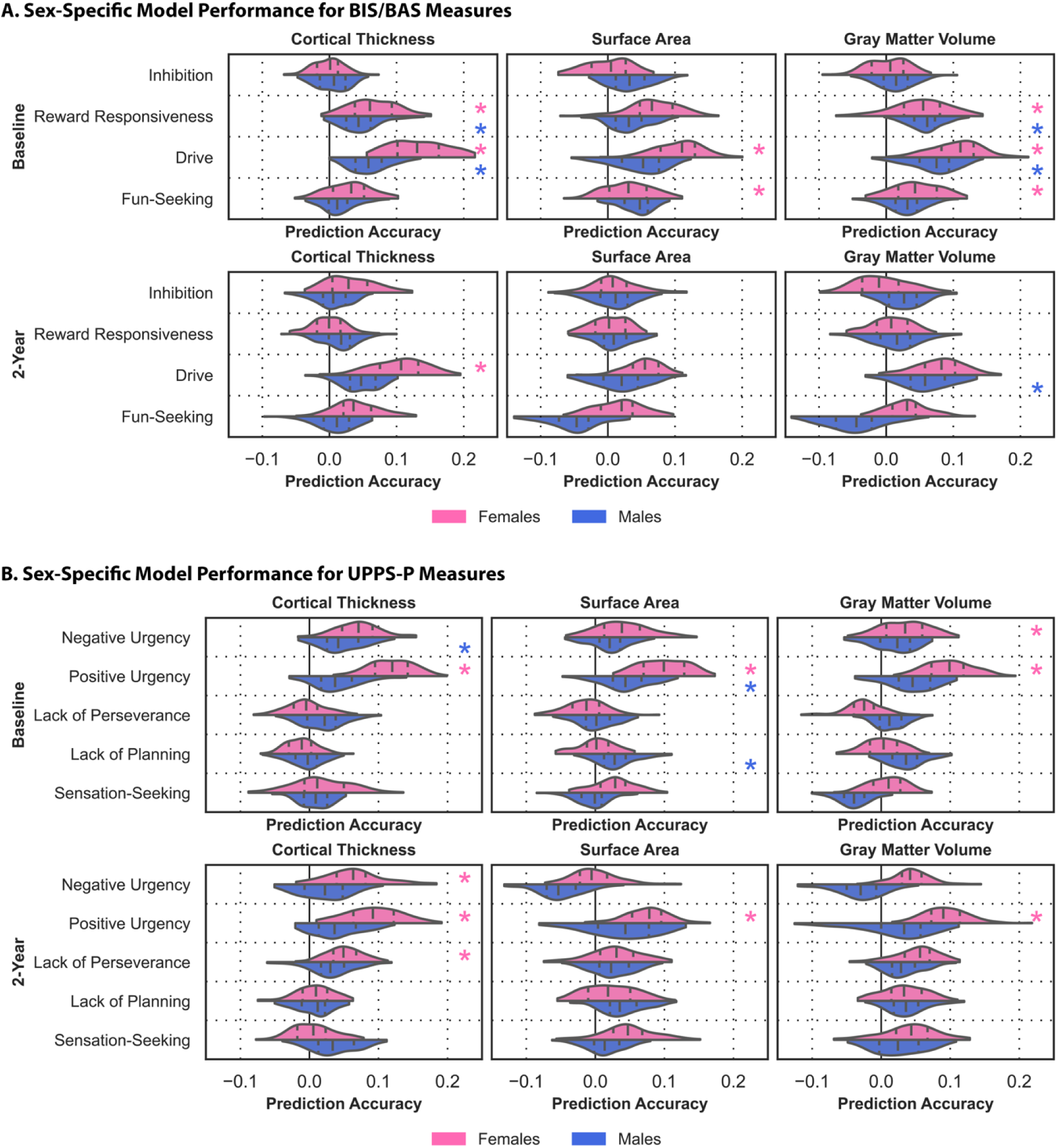
Sex influences associations between neuroanatomy and impulsivity. Prediction accuracies (correlation between observed and prediction values) for sex-specific models trained to predict behavioral measures from the BIS/BAS (A) and UPPS-P (B) scales. Results for female-specific (pink) and male-specific (blue) models based on CT (left), SA (center), and GMV (right) at baseline (top) and two-year follow-up (bottom) are shown. The shape of the violins indicates the distribution of values, the dashed lines indicate the median, and the dotted lines indicate the interquartile range. Asterisks indicate the model captured significant associations.

<u>CT:</u> Sex-specific models based on CT accurately predicted reward-responsiveness (r_female_=0.065, p_female_=0.002; r_male_=0.046, p_male_=0.008) and drive (r_female_=0.133, p_female_<0.001; r_male_=0.060, p_male_=0.004) in both sexes, negative urgency (r=0.048, p=0.015) in males, and positive urgency (r=0.119, p<0.001) in females at baseline; and drive (r=104, p=0.028), negative urgency (r=0.065, p=0.005), positive urgency (r=0.093, p<0.001), and lack of perseverance (r=0.051, p=0.038) in females at two-year follow-up.

<u>SA:</u> Sex-specific models based on SA accurately predicted positive urgency (r_female_=0.099, p_female_<0.001; r_male_=0.046, p_male_<0.005) in both sexes, drive (r=0.106, p<0.001) and fun-seeking (r=0.045 p=0.006) in females, and lack of planning (r=0.031, p=0.008) in males at baseline; and positive urgency in females (r=0.075, p=0.005) at two-year follow-up.

<u>GMV:</u> Sex-specific models based on GMV accurately predicted reward-responsiveness (r_female_=0.052, p_female_=0.028; r_male_=0.059, p_male_=0.020) and drive (r_female_=0.105 p_female_<0.001; r_male_=0.072, p_male_<0.001) in both sexes, fun-seeking (r=0.045, p=0.006), and negative urgency (r=0.034, p=0.040) in females at baseline; and drive (r=0.060, p<0.001) in males and positive urgency in females (r=0.094, p=0.005) at two-year follow-up.

Changes in impulsivity are part of normative development and, at their extremes, are linked to the development of mental illnesses later in life^60,61^. We show that individual variations in neuroanatomy can predict impulsivity in youth, and many of these relationships are stable across development. While certain measures can be predicted in both sexes, others yield significant results only in females, which may be due to the reliability of measures across sexes, reporting biases, and data quality, among other factors.

### Impulsivity maps onto shared and distinct brain regions

We derived the feature importance maps from the models and computed cosine similarities to evaluate overlap, focusing on models that captured significant associations. These results, presented in their entirety in **Figure 3A**, indicated that models predicting measures from the same scale captured largely overlapping associations. As an example, CT features associated with positive and negative urgency were highly similar at both time points (cosine similarity, s_baseline_=0.69, s_two-year_=0.77). Further, while some models predicting measures from different scales were similar, others were orthogonal or opposite. For example, baseline CT features associated with reward-responsiveness and drive were dissimilar from those associated with lack of planning and sensation seeking (−0.60≤s≤-0.29), but similar to those associated with positive urgency (0.62_≤_s_≤_0.64). These observed patterns were generally consistent across time points for CT and SA, but less so for GMV.

**Figure 3:**
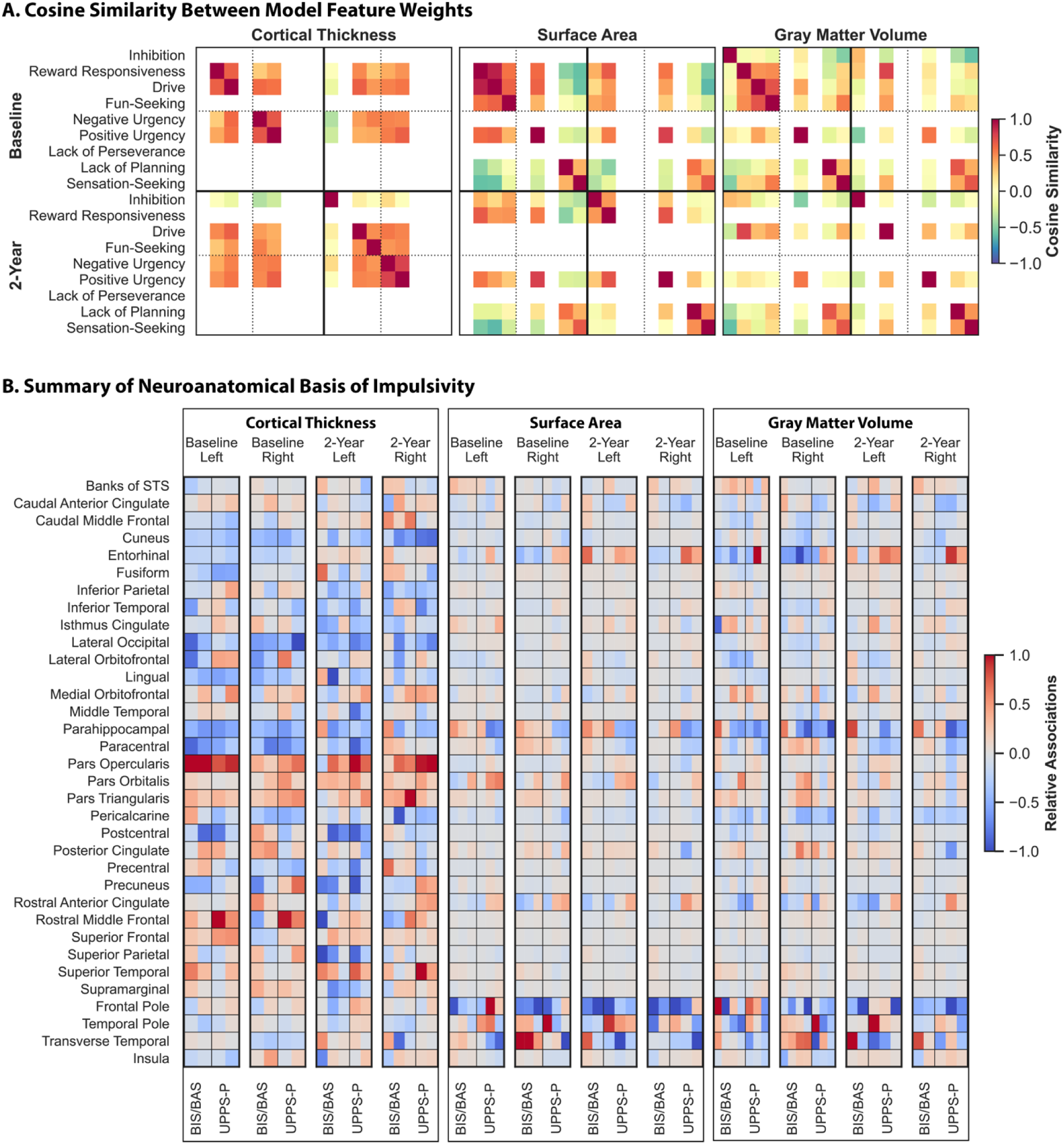
Shared and unique neuroanatomical features are linked to impulsivity. Cosine similarities between the Haufe-transformed regional feature weights from models trained to predict impulsivity (A). Results for models based on CT (left), SA (center), and GMV (right) that captured significant associations are shown. Warmer colors indicate greater similarity, cooler colors indicate a greater dissimilarity. Rows and columns corresponding to models that did not capture significant associations are left blank. Relative regional associations between neuroanatomy and impulsivity derived from the models based on CT (left), SA (center), and GMV (right) (B). Left and right hemisphere are shown as denoted by the top, x-axis labels. Warmer colors indicate a stronger positive association, cooler colors indicate a stronger negative association. To facilitate visualization, association values for each set of models were divided by the maximum value for that model. Results are only shown for models that captured significant associations, as outlined below (ordered left to right), and measures from the scales are separated by vertical lines. CT, Baseline: Reward Responsiveness, Drive, Negative Urgency, Positive Urgency CT, Two-Year: Inhibition, Drive, Fun-Seeking, Negative Urgency, Positive Urgency SA, Baseline: Reward Responsiveness, Drive, Fun-Seeking, Positive Urgency, Lack of Planning, Sensation-Seeking SA, Two--Year: Inhibition, Reward Responsiveness, Positive Urgency, Lack of Planning, Sensation-Seeking GMV, Baseline: Inhibition, Reward Responsiveness, Drive, Fun-Seeking, Positive Urgency, Lack of Planning, Sensation-Seeking GMV, Two-Year: Inhibition, Drive, Fun-Seeking, Positive Urgency, Lack of Planning, Sensation-Seeking

We also evaluated the similarities in the neuroanatomical features associated with impulsivity measures across the sexes, focusing on the five pairs of models that yielded significant results in both sexes at baseline. GMV features associated with reward-responsiveness at baseline were quite similar across the sexes (s=0.53), along with SA features associated with positive urgency (s=0.53). However, other associations were considerably different across the sexes. Models based on CT to predict reward-responsiveness and drive captured distinct associations in males and females (s_reward-responsiveness_=0.24, s_drive_=0.37), as well as those based on GMV to predict drive (s=0.36).

These analyses reveal that impulsivity maps onto shared and distinct neuroanatomical features. Across the entire sample, reward sensitivity and urgency share a common neuroanatomical basis that is distinct from the neural substrates of lack of planning and sensation-seeking. These findings suggest that the BIS/BAS and UPPS-P scales, though separable constructs, share some neuroanatomical features while also exhibiting distinct features. Importantly, some of these associations differ across the sexes, suggesting the presence of sex-specific neuroanatomical substrates.

### Neuroanatomical basis of impulsivity varies across imaging modalities

CT features associated with impulsivity were widespread, while SA and GMV features overlapped and were more localized (full results shown in **Figure 3B**).

<u>CT:</u> Impulsivity was broadly negatively associated with CT in bilateral cuneus and lateral occipital regions (i.e., youth who were more impulsive had less CT in these areas relative to those who were less impulsive) and was positively associated with CT in the bilateral inferior frontal gyrus, particularly in the pars opercularis, superior frontal, and superior temporal regions (**Figure 4A-D** for 2 representative BIS/BAS and UPPS-P measures at each time point, **Figure S7** for all other measures). Impulsivity was also negatively associated with CT in bilateral entorhinal, lingual, parahippocampal, and paracentral regions at baseline, but these associations were less consistent at two-year follow-up. Other regions exhibited measure- and hemisphere-specific relationships. For example, impulsivity was broadly negatively associated with CT in the left postcentral and precuneus regions, but BIS/BAS measures were positively associated with CT in the right postcentral region while UPPS-P measures were positively associated with CT in right precuneus at baseline.

**Figure 4.**
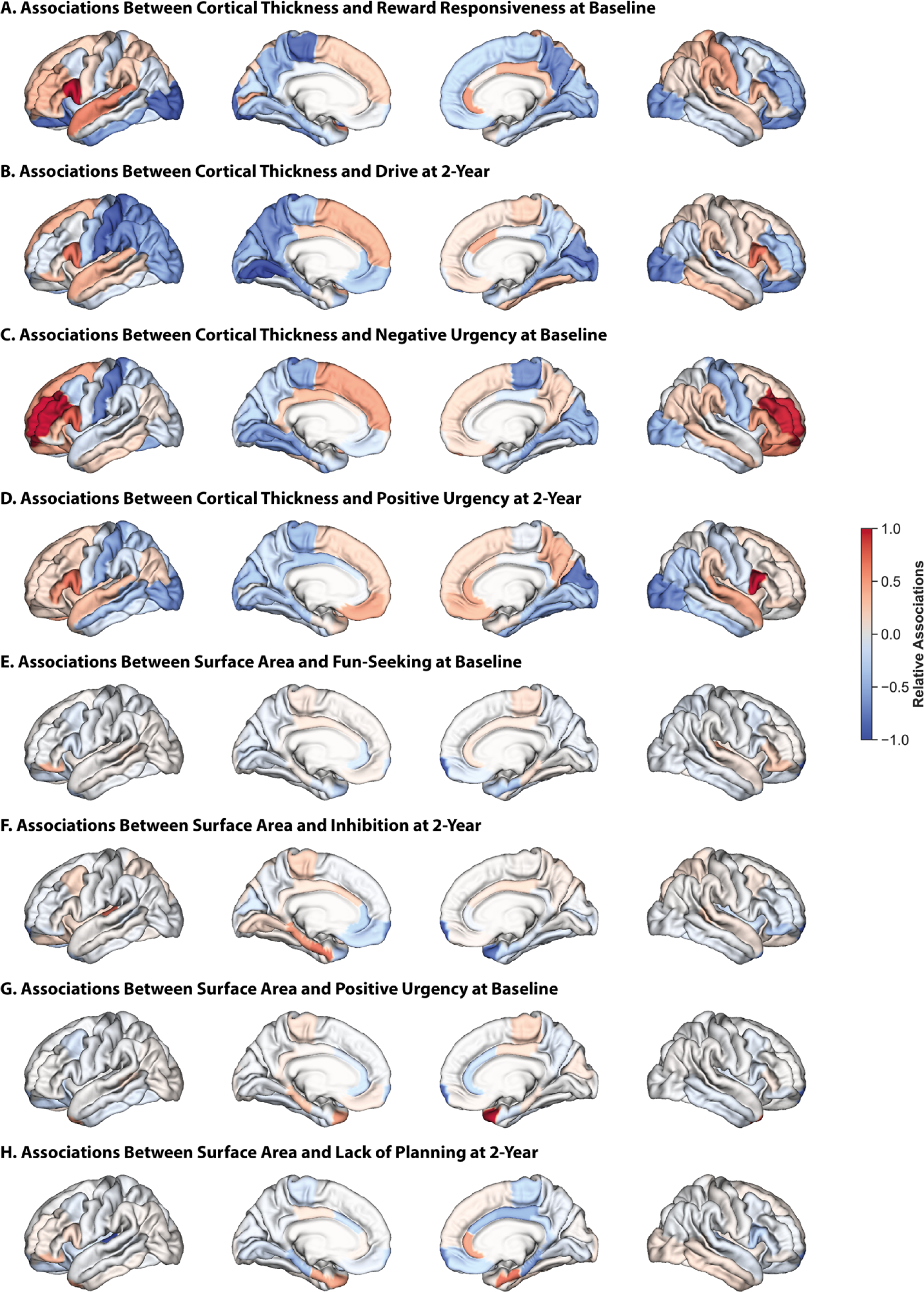
Associations between CT and impulsivity are dispersed throughout the cortex, while those between SA (and gray matter) and impulsivity are localized. Relative regional associations from models trained on CT data to predict reward-responsiveness at baseline (A), drive at two-year follow-up (B), negative urgency at baseline (C), and positive urgency at two-year follow-up (D). Relative regional associations from models trained on SA data to predict fun-seeking at baseline (E), inhibition at two-year follow-up (F), positive urgency at baseline (G), lack of planning at two-year follow-up (H). Lateral (outer) and medial (inner) surfaces for left (left) and right (right) hemispheres are shown. Warmer colors indicate a stronger positive association, cooler colors indicate a stronger negative association. To facilitate visualization, association values for each set of models were divided by the maximum value for that model.

<u>SA:</u> Impulsivity was broadly negatively associated with SA in bilateral frontal poles at both time points (**Figure 4E-H** for 2 representative BIS/BAS and UPPS-P measures at each time point, **Figure S8** for all other measures). Positive urgency was also positively associated with SA in bilateral temporal poles, but associations with temporal pole regions were more nuanced for other measures. Certain regions also exhibited opposite associations. For example, BIS/BAS measures were positively associated with SA in the transverse temporal and parahippocampal regions and negatively associated with SA in the rostral anterior cingulate and entorhinal regions, while the opposite was true for UPPS-P measures.

<u>GMV:</u> Associations between GMV and impulsivity largely paralleled those with SA, although they were less pronounced and included a few exceptions (**Figure S9** for 2 representative BIS/BAS and UPPS-P measures at each time point, **Figure S10** for all other measures). Impulsivity was broadly negatively associated with GMV in the pericalcarine region, and inhibition was positively associated with frontal pole volumes but negatively associated with isthmus cingulate volumes.

These findings highlight the presence of multivariate relationships between neuroanatomy and impulsivity. While some associations are stable across the behavioral scales and time points, others are less consistent. These results suggest a distributed network of brain regions encode individual differences in impulsive behaviors, and these relationships are dynamic.

### There are sex differences in the neuroanatomical basis of impulsivity

We assessed sex differences in the associations between neuroanatomy and impulsivity, focusing on the models that were significant in both sexes. Across both sexes, BAS measures were negatively associated with CT in the entorhinal, lateral occipital, lateral orbitofrontal, and parahippocampal regions (**Figure 5**). In females, they were also positively associated with CT in bilateral inferior frontal gyrus and, to a lesser extent, in the superior frontal gyrus, while in males, these relationships were present in the left hemisphere, but the opposite relationships were present in the right hemisphere. Further, although these measures were negatively associated with CT in the lingual, paracentral, and precuneus regions in both sexes, these relationships were stronger in the lingual regions in females and in the paracentral and precuneus regions in males.

**Figure 5:**
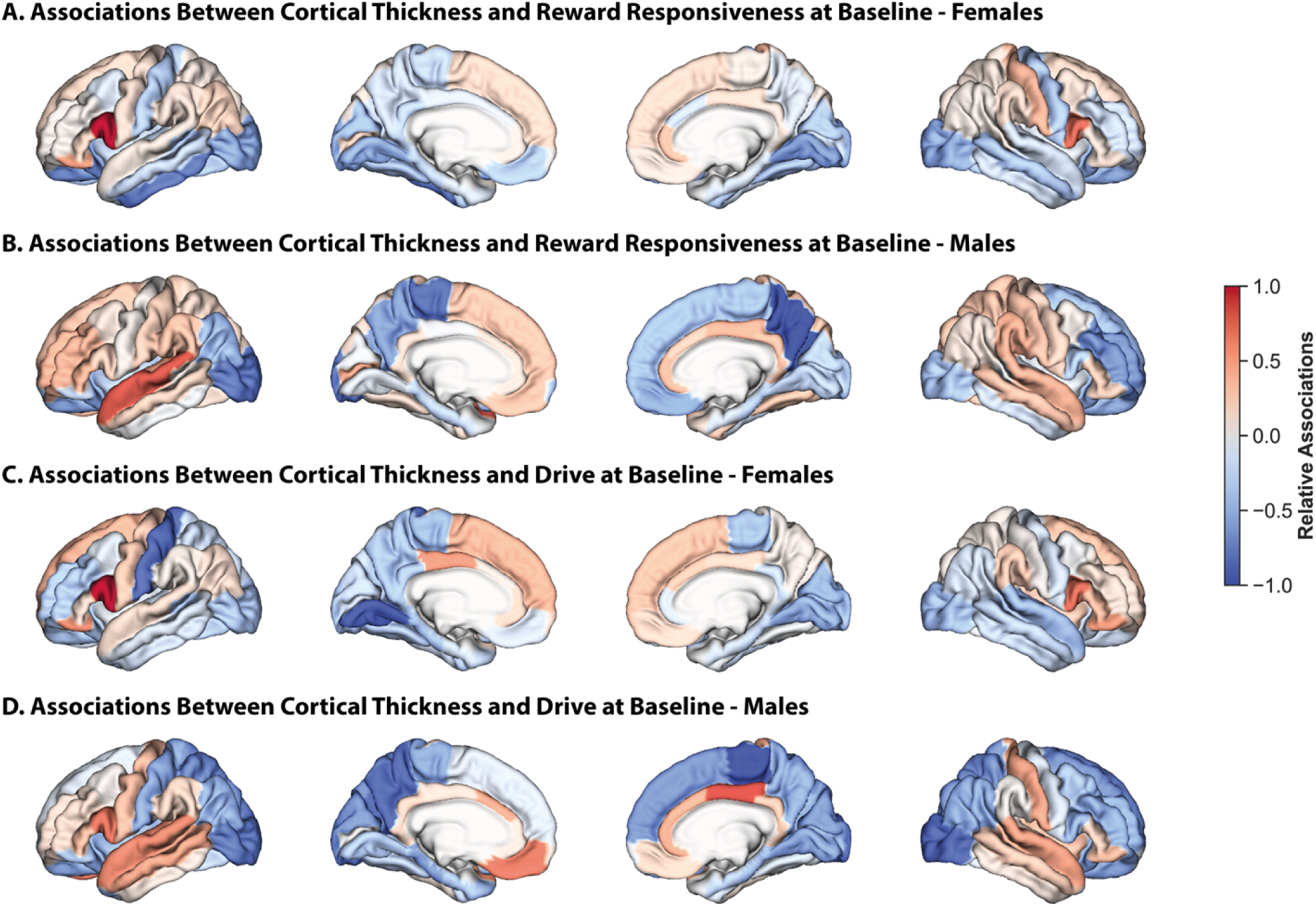
There are sex differences in the associations between CT and reward sensitivity. Relative regional associations from models trained on CT data to predict reward-responsiveness at baseline in females (A) and males (B), and drive at baseline in females (C) and males (D). Lateral (outer) and medial (inner) surfaces for left (left) and right (right) hemispheres are shown. Warmer colors indicate a stronger positive association, cooler colors indicate a stronger negative association. To facilitate visualization, association values for each set of models were divided by the maximum value for that model.

Relationships between GMV and reward-responsiveness at baseline were largely shared across the sexes, but those between GMV and drive were considerably different, although the differences were predominantly in strength rather than directionality (**Figure S11**). As an example, GMV in the right posterior cingulate and right pars opercularis exhibited stronger relative positive associations with drive in females than in males. Results from all other sex-specific significant models generally resembled the results from the sex-independent models (**Figures S12-14)**.

These findings suggest that while many of these relationships are shared across the sexes, there are also important differences in the neuroanatomical underpinnings of impulsivity. These sex-specific relationships may, in part, explain observed sex differences in impulsive behaviors and vulnerability to impulsivity-related psychiatric illness.

## Discussion

Examining multiple facets of impulsivity and neuroanatomical features, we demonstrate that individual variability in neuroanatomy in a large cohort of youth is associated with impulsivity across facets and features, and these relationships, to some extent, differ between sexes. We show that impulsivity measures map onto shared and distinct brain regions across neuroanatomical features. Some relationships overlap across the impulsivity measures and across neuroanatomical features, are consistent across the sexes, and remain stable during development, while others vary across facets, features, sexes, and time points. These results shed light on how individual differences in neuroanatomy throughout development contribute to the diverse expressions of impulsivity in youth. These findings also suggest that impulsivity maps onto specific patterns of neuroanatomy that, alongside other risk factors, could help identify youth at risk for impulsivity-related disorders.

Impulsivity is, in part, driven by individual differences in corticolimbic, corticostriatal and motor-sensory circuits^8,13–16,62^. Recent advances in brain-based predictive modeling allow us to examine whole-brain multivariate relationships^31^, unlike traditional univariate analyses focused on individual regions. In recent years, a few studies have used this approach to investigate the neuroanatomical basis of impulsivity, although they have generally focused on specific neuroimaging features and impulsivity measures, ignored sex effects, and considered these relationships at a single time point^8,52,63,64^. Using a multivariate approach, we broadly replicate univariate findings and demonstrate that impulsivity maps onto a dispersed set of cortical regions. While some relationships are shared across facets of impulsivity, others are distinct. One particularly stable relationship appears to be in the pars opercularis, where CT is positively linked to impulsivity measures from both scales at both time points. These results align with those from prior neuroanatomical, functional, and electrophysiological studies showing that the pars opercularis plays an important role in impulse control^65^ across motor^66–68^ and speech^69,70^ domains.

Impulsivity follows a non-linear trend, increasing during childhood and adolescence and decreasing throughout adulthood^71–73^. This trajectory mirrors developmental changes observed in neuroanatomy. During adolescence and early adulthood, significant maturation occurs in the prefrontal cortex, a region critical for emotional regulation and impulse control^20–22^. This maturation involves synaptic pruning and increased white matter connectivity to refine neural circuits, leading to improved cognitive control and decreased impulsivity^22^. Our results show that the neuroanatomical basis of impulsivity is not static, potentially reflecting these broader developmental changes. While some regions show consistent relationships, others demonstrate changes from baseline to two-year follow-up, suggesting that changes in impulsivity may be driven by shifts in the underlying neuroanatomical associations. Further, deviations from typical developmental trajectories in neuroanatomy may underlie impulsivity-related deficits.

Research on sex differences in impulsivity has produced mixed findings^36,74^. The most consistent finding is that females exhibit greater inhibition and males exhibit greater sensation-seeking^36^. Activation-related impulsivity is comparable across the sexes^36^, though differences have been reported for specific rewards^75^. One study examining relationships between CT and a single global measure of impulsivity in the ABCD cohort reported significant associations in males but not the entire sample^76^. A separate study exploring the volumetric correlates of impulsivity in the same cohort found that lack of premeditation and sensation seeking were related to larger volumes in many cortical and subcortical regions, while positive urgency was related to smaller volumes in those same regions^77^. They also found that many of the relationships were stronger in females. These studies highlight the need for sex-specific investigations. Our analyses build on this work using a multivariate machine learning approach and show that while some brain-impulsivity associations are consistent across the sexes, others are not. In some cases, the same regions even exhibit opposite relationships across the sexes.

There are several limitations to this work. First, we used a single dataset. Although participants reflect different demographic groups, income levels, and living environments, our findings may be limited in generalizability^78,79^. To maximize the robustness of our work, we included all participants with complete imaging and impulsivity data and used a cross-validated predictive modeling framework known to yield reliable results^50^. Second, we only considered binary sex due to data availability and thus were not able to assess these relationships in intersex or other non-binary populations. We also did not consider the effects of gender, which influences neurobiology^46^ and behavior^80^. Third, the brain continues to develop throughout adolescence, with females and males reaching developmental milestones at different times^81^. Here, we used data from two time points but did not include older ages due to data availability. While some neuroanatomical associations with impulsivity were stable, others shifted over time. As a result, the associations we report may continue to shift throughout development. Our work serves as an empirical baseline from which theses trajectories may be tracked in later waves of the ABCD study. Finally, we examined the neural basis of impulsivity and explored sex differences in these relationships. However, we did not consider the effects of other biological or environmental factors (e.g., genetics, pubertal maturation, urbanicity)^82–85^. Future analyses within global open-access datasets that consider the effects of additional biological and environmental factors can address these limitations and provide additional evidence to confirm (or refute) these findings.

Increases in impulsivity are a typical part of development, and, when significant, may be linked to risk for psychiatric illness. Understanding the neuroanatomical basis of impulsivity paves the way for development of more effective early interventions grounded in neurobiological mechanism to prevent psychiatric illness. These findings highlight how neuroanatomy underlies the diverse expressions of impulsivity throughout development and may represent potential markers of psychiatric risk. In addition, this work emphasizes the importance of conducting sex-disaggregated analyses when examining brain-behavior relationships.

## Data Availability Statement

Data used in the preparation of this article were obtained from the Adolescent Brain Cognitive Development^SM^ (ABCD) Study (https://abcdstudy.org), held in the NIMH Data Archive (NDA). This is a multisite, longitudinal study designed to recruit more than 10,000 children aged 9-10 and follow them over 10 years into early adulthood. The ABCD Study® is supported by the National Institutes of Health and additional federal partners under award numbers U01DA041048, U01DA050989, U01DA051016, U01DA041022, U01DA051018, U01DA051037, U01DA050987, U01DA041174, U01DA041106, U01DA041117, U01DA041028, U01DA041134, U01DA050988, U01DA051039, U01DA041156, U01DA041025, U01DA041120, U01DA051038, U01DA041148, U01DA041093, U01DA041089, U24DA041123, U24DA041147. A full list of supporters is available at https://abcdstudy.org/federal-partners.html. A listing of participating sites and a complete listing of the study investigators can be found at https://abcdstudy.org/consortium_members/. ABCD consortium investigators designed and implemented the study and/or provided data but did not necessarily participate in the analysis or writing of this report. This manuscript reflects the views of the authors and may not reflect the opinions or views of the NIH or ABCD consortium investigators.

