## Supplemental Materials for "Neuroanatomy Reflects Individual Variability in Impulsivity in Youth"

**Introduction**

Heightened impulsivity is a core feature of many psychiatric disorders^1^. While certain disorders are associated with broad impulsivity, others are linked to specific impairments. Individuals with a history of interpersonal childhood trauma^2^, major depressive disorder^3,4^, bipolar disorder^3,4^, anorexia nervosa^5^, or social anxiety disorder^6^ exhibit heightened levels of behavioral inhibition relative to controls. Individuals with bipolar disorder additionally exhibit greater activation relative to controls^3,7^, while the opposite is observed in those with major depressive disorder^6^, social anxiety disorder^6^, or schizophrenia^7^. Further, individuals with borderline personality disorder^8^ substance use disorder^9^, or attention deficit hyperactivity disorder^10^ exhibit higher levels of impulsivity across the UPPS-P measures compared to controls. In contrast, individuals with generalized anxiety disorder specifically exhibit higher levels of negative urgency and lack of premeditation^11^. Finally, the onset of self-harm behaviors is linked to heightened sensation-seeking, while the maintenance of these behaviors is linked to heightened lack of premeditation^12^. Impulsivity also affects physical health, with higher levels associated with various illnesses across multiple biological systems, including endocrine/metabolic, neurological, circulatory, respiratory, and digestive^13^.

**Methods**

**Dataset**

We used ABCD Study data from the ABCD 5.1 release. The research protocol for the dataset was reviewed and approved by a central Institutional Review Board (IRB) at the University of California, San Diego, and, in some cases, by individual site IRBs. Parents or guardians provided written informed consent, and children assented before participation. We excluded participants with imaging data that was missing, as well as those not recommended for inclusion or with incidental MRI findings, as recommended in the ABCD Release Notes (<https://wiki.abcdstudy.org/release-notes/imaging/quality-control.html>). We also excluded participants with incomplete self-report trait data. Finally, we excluded related participants such that a single family member was included in the sample and others were dropped at random. We included 9,099 participants at baseline and 6,432 participants at two-year follow-up with complete and useable imaging and impulsivity data (see **Figures S1-S2** for our participant inclusion pipeline and **Table S1** for participant demographic data).

**Impulsivity**

BIS/BAS: We derived participant scores from the Behavioral Inhibition/ Activation System (BIS/BAS) scale. The BIS/BAS scale is a reliable and well-validated 20-item self-report questionnaire used to measure individual differences in the behavioral inhibition and activation systems^14-17^. The behavioral inhibition system reflects motivation to avoid aversive outcomes, and the behavioral activation system reflects motivation to approach goal-oriented outcomes. There is one BIS-related scale (inhibition), and three BAS-related scales (reward responsiveness, drive, and fun-seeking). Inhibition reflects sensitivity to punishment and is the sum of seven items. Reward responsiveness reflects reward anticipation, reward response, and reward satiation, and is the sum of four items. Drive reflects persistent pursuit of goals and is the sum of four items. Fun-seeking reflects desire to approach potential rewards impulsively and is the sum of five items. Participants respond to each item using a four-point Likert scale (ranging from 0 to 3), where higher values indicate higher levels of a given trait.

UPPS-P: We derived participant scores from the Modified Urgency, (lack of) Planning (or Premeditation), (lack of) Perseverance, Sensation-Seeking, and Positive Urgency (UPPS-P^18^) Short Version scale. The UPPS-P scale is a reliable and well-validated 20-item self-report questionnaire used to measure five distinct impulsive personality traits: negative urgency, positive urgency, lack of planning, lack of perseverance, and sensation-seeking^19-21^. Negative urgency is the tendency to act impulsively to negative emotions. Positive urgency is tendency to act impulsively to positive emotions. Lack of perseverance is the tendency to give up or not complete tasks. Lack of planning is the tendency to act without considering the consequences. Sensation-seeking is the tendency to pursue exciting or novel activities. Scores for each of the 5 traits are based on the sum of four items each. Participants respond to each item using a four-point Likert scale (ranging from 1 to 4) where higher values indicate higher levels of a given trait.

**Predictive Modelling**

For each set of models, we first randomly shuffled and split the data into 100 distinct training and test sets (at approximately a 4:1 ratio) without replacement. We accounted for potential variability introduced by imaging site by placing all participants from a given site either in the train set or the test set but not split across the two. Within each training set, we optimized the regularization hyperparameter using three-fold cross-validation, again ensuring participants from a given site were not split across folds. Once optimized, we trained the model on the entire training set using the optimized hyperparameter and evaluated model performance on the corresponding test set based on prediction accuracy (i.e., correlation between observed and predicted values), as in prior work^22-25^. We repeated this process for all training-test splits to obtain a distribution of prediction accuracy for each set of models. We evaluated model significance by comparing these distributions to corresponding null distributions. We generated null models by randomly permuting the output variable within each site and then using those data to train and test a model using a randomly selected regularization hyperparameter from the set of optimized hyperparameters for the original model. This was repeated 1000 times to generate a null distribution. We obtained the p-values for model performance by calculating the proportion of null models with prediction accuracies greater than or equal to the corresponding original distribution. We corrected the p-values for multiple comparisons within the behavioral scales the Benjamini-Hochberg False Discovery Rate (q=0.05) procedure^26^.

Of note, the inclusion of covariates in brain-based predictive models can mask crucial interactions and non-linear relationships between brain features and the outcome variable^27,28^. Covariate adjustment generally assumes a linear and independent relationship between the covariate and the outcome, which may not hold true in complex biological systems. By not including covariates, we allow the model to capture the full range of variation in the neuroimaging data, including variance that might be shared with typical covariates, thus potentially revealing more robust and biologically relevant predictive patterns. Therefore, we did not include any covariates in these analyses.

**Feature Weights**

Cosine similarity is calculated using the dot product of two vectors divided by the product of their magnitudes. Cosine similarity values range from -1 to 1, where 1 indicates complete similarity, 0 indicates orthogonality or no similarity, and -1 indicates complete dissimilarity.

**Results**

Within the measures derived from the BIS/BAS scale, youths exhibited significantly lower levels of all four behaviors at the two-year follow-up (Mann Whitney U statistic, U_inhibition_=3.3x10^7^; U_reward responsiveness_=3.5x10^7^; U_drive_=3.2x10^7^; U_fun-seeking_=3.6x10^7^; all p_FDR_<0.001; **Figure S5A**). Within the measures derived from the UPPS-P scale, youths exhibited lower levels of negative urgency (U=3.4x10^7^, p_FDR_<0.001), positive urgency (U=3.2x10^7^, p_FDR_<0.001), lack of planning (U=2.8x10^7^, p_FDR_=0.004), and sensation-seeking (U=3.1x10^7^, p_FDR_<0.001) at the two-year follow-up, but similar levels of lack of perseverance (U=3.0x10^7^, p_FDR_=0.241; **Figure S5B**).

Sex differences were present in all behavioral measures (**Figure S6;** all p_FDR_ <0.001 unless indicated otherwise). Females exhibited significantly higher levels of inhibition (U_baseline_=1.1x10^7^; U_2-year_=6.0x10^6^) but lower levels of reward responsiveness (U_baseline_=9.7x10^6^; U_2-year_=5.0x10^6^, p_FDR_=0.03), drive (U_baseline_=9.2x10^6^; U_2-year_=4.6x10^6^), and fun-seeking (U_baseline_=9.6x10^6^; U_2-year_=4.7x10^6^). Females also exhibited significantly lower levels of negative urgency (U_baseline_=9.2x10^6^; U_2-year_=4.6x10^6^), positive urgency (U_baseline_=9.3x10^6^; U_2-year_=4.5x10^6^), lack of perseverance (U_baseline_=9.6x10^6^; U_2-year_=4.9x10^6^), lack of planning (U_baseline_=9.0x10^6^; U_2-year_=4.5x10^6^), and sensation-seeking (U_baseline_=9.6x10^6^; U_2-year_=4.9x10^6^).

Across the entire sample, reward responsiveness, drive, and fun-seeking were moderately correlated (0.44≤r_baseline_≤0.46; 0.44 ≤r_2-year_≤0.46), negative and positive urgency were moderately correlated (r_baseline_=0.49; r_2-year_=0.53), and lack of perseverance and planning were moderately correlated (r_baseline_=0.45; r_2-year_=0.47). Across the behavioral scales, fun-seeking and sensation-seeking were moderately correlated (r_2-year_=0.41) and all other measures were weakly correlated (|r|<0.40), if at all (**Figure S5C**). Correlation values with an absolute magnitude below 0.02 were not statistically significant after FDR correction, but all others were significant (p_FDR_ <0.050). Similar patterns were also present within each sex (**Figure S6**).

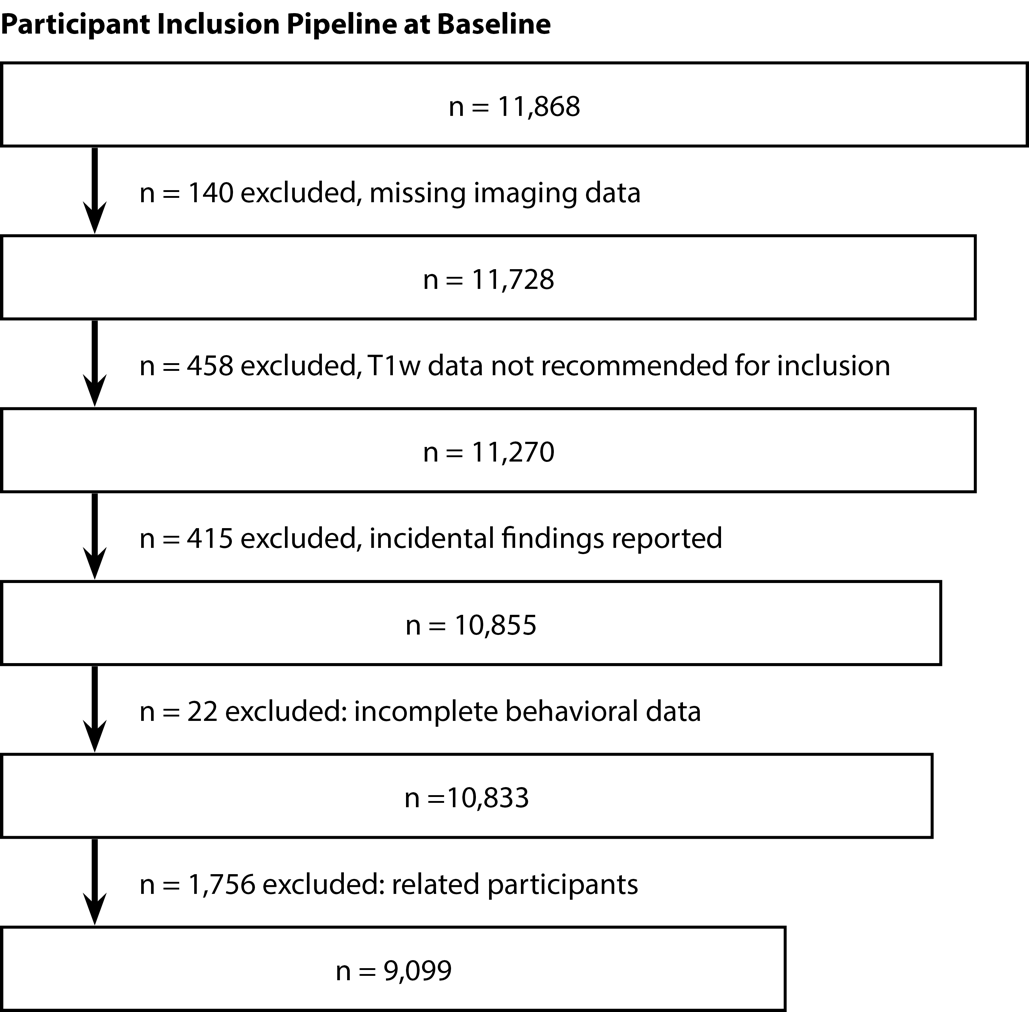

**Figure S1**: **Participant inclusion/exclusion workflow at baseline.**
Overview of the process used to include participants in these analyses at baseline.

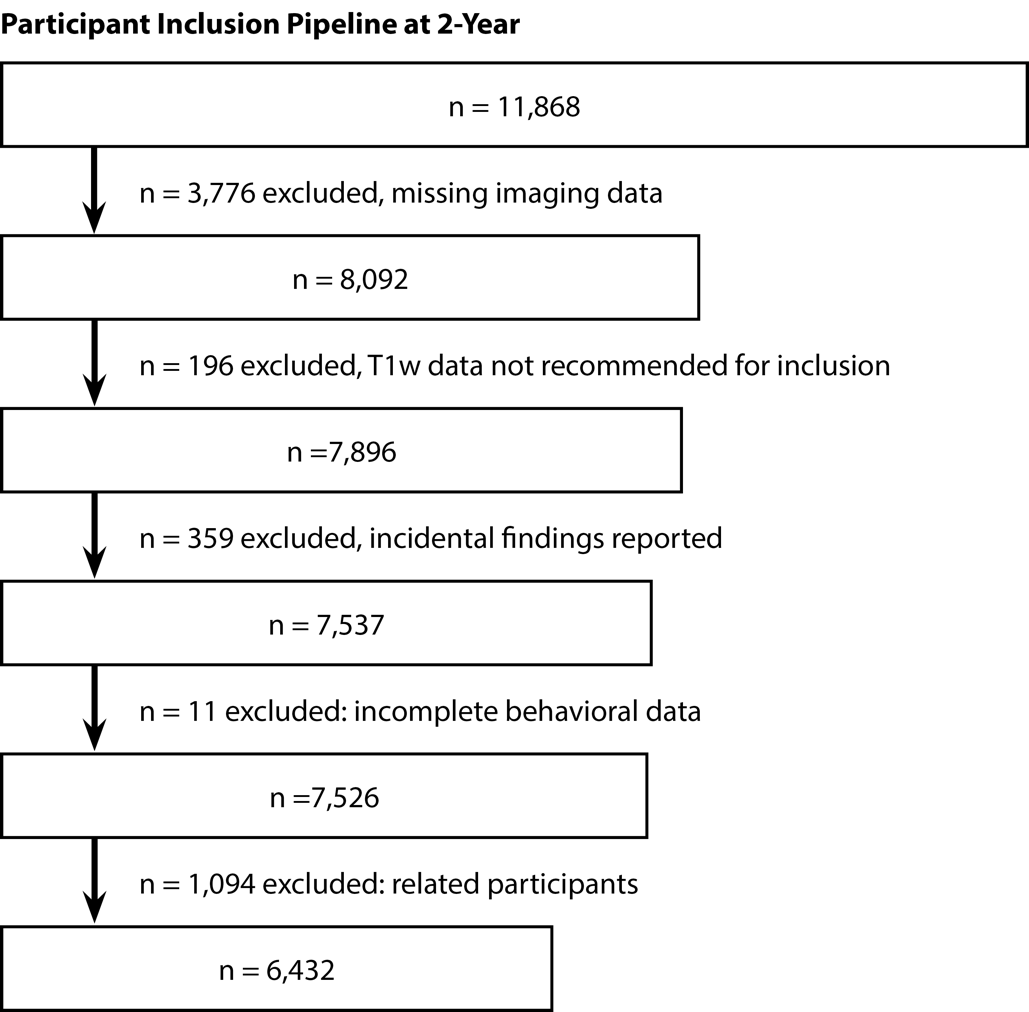

**Figure S2:** **Participant inclusion/exclusion workflow at 2-year follow-up.**
Overview of the process used to include participants in these analyses at 2-year follow-up.

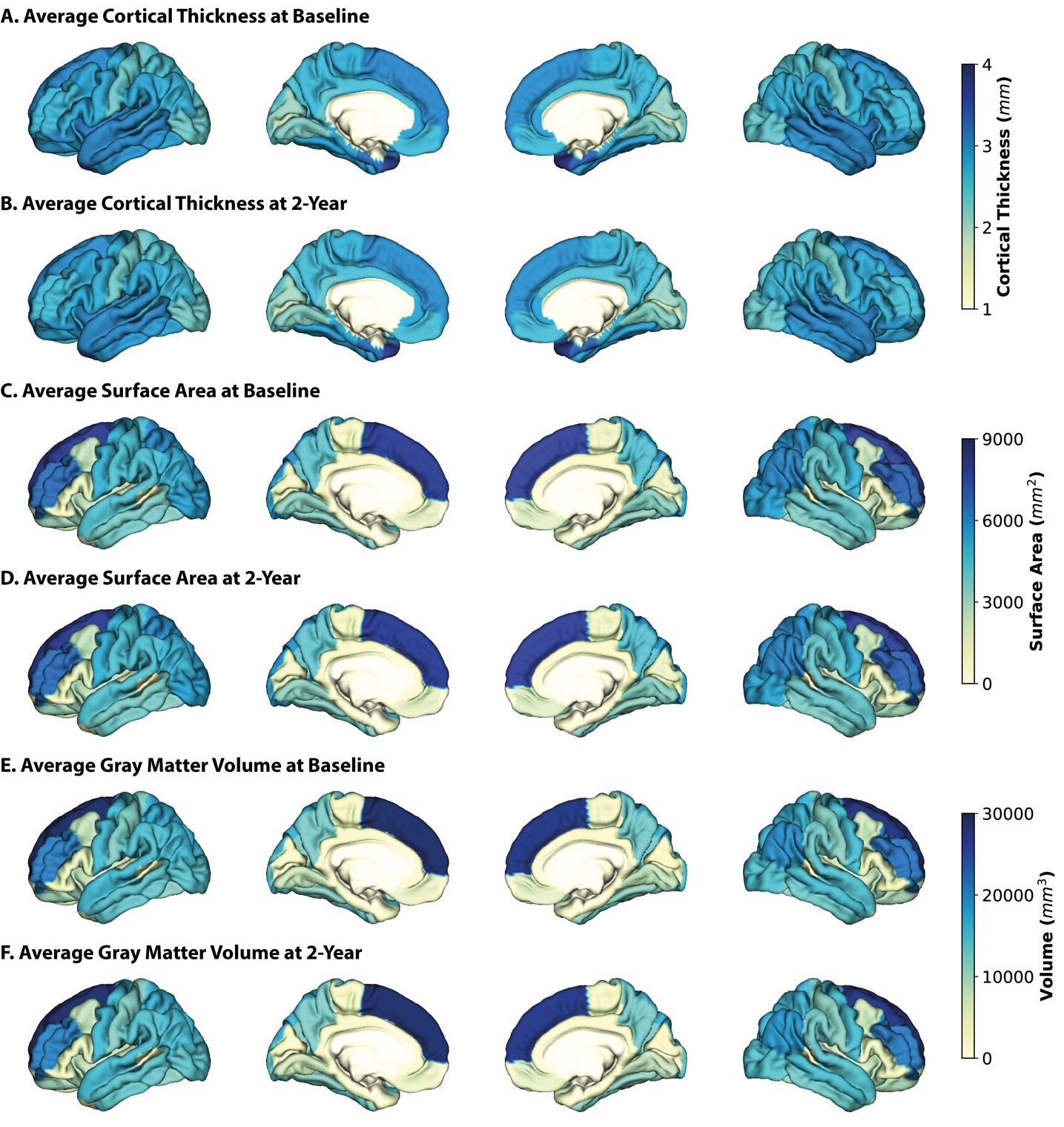

**Figure S3: Average neuroanatomical measures at baseline and 2-year follow-up.**
Average cortical thickness at baseline (A) and 2-year follow-up (B), average surface area at baseline (C) and 2-year follow-up (D), and average gray matter volume at baseline (E) and 2-year follow-up (F). Lateral and medial surfaces for left and right hemispheres are shown.

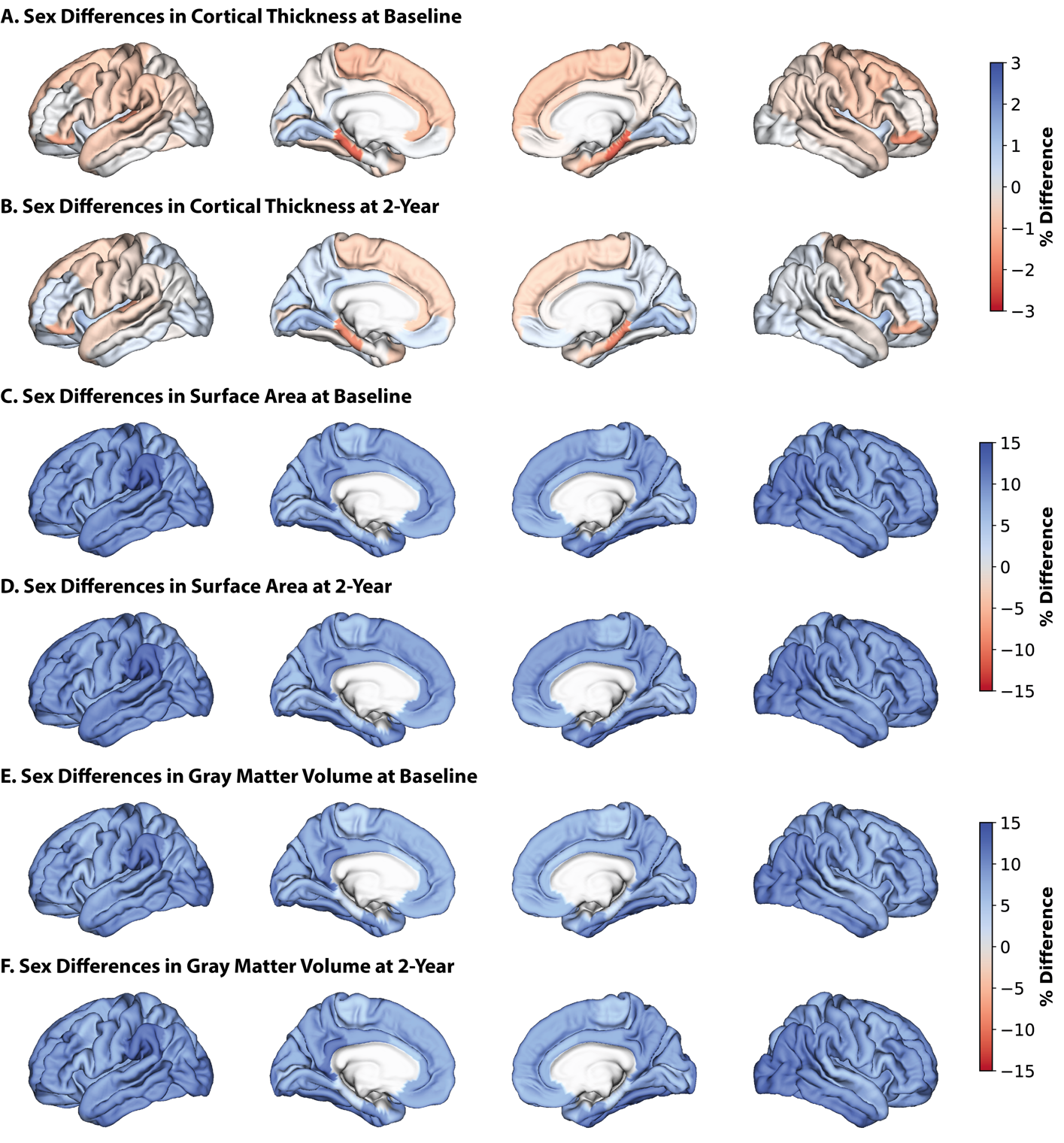

**Figure S4: Sex differences in neuroanatomical measures at baseline.**
Percent difference between the sexes in cortical thickness at baseline (A) and 2-year follow-up (B), average surface area at baseline (C) and 2-year follow-up (D), and average gray matter volume at baseline (E) and 2-year follow-up (F). Percent differences were calculated by subtracting regional measures in females from regional measures in males and dividing by regional measures in females and then multiplying by 100. Warmer colors indicate a higher thickness/area/volume in females relative to males, cooler colors indicate a higher thickness/area/volume in males. Lateral and medial surfaces for left and right hemispheres are shown.

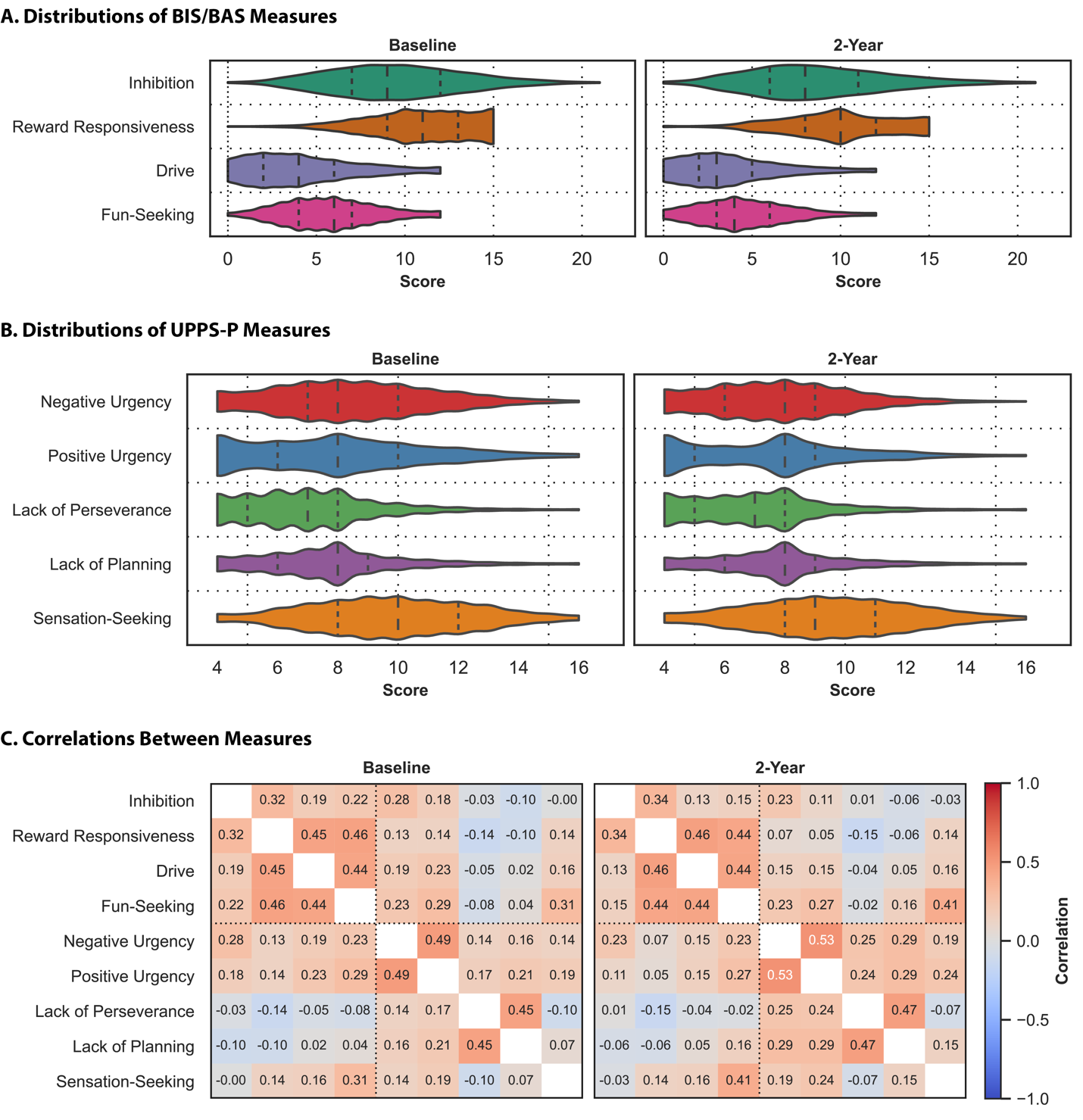

**Figure S5**: **Participants exhibit varying levels of impulsivity, and the measures reflect independent constructs.**
Distribution of behavioral measures from the BIS/BAS (A) and UPPS-P (B) scales. The shape of the violins indicates the distribution of values, the dashed lines indicate the median, and the dotted lines indicate the interquartile range. Correlations between behavioral measures (C). Baseline (left) and two-year follow-up (right) data are shown. Warmer colors indicate a stronger positive correlation, cooler colors indicate a stronger negative correlation.

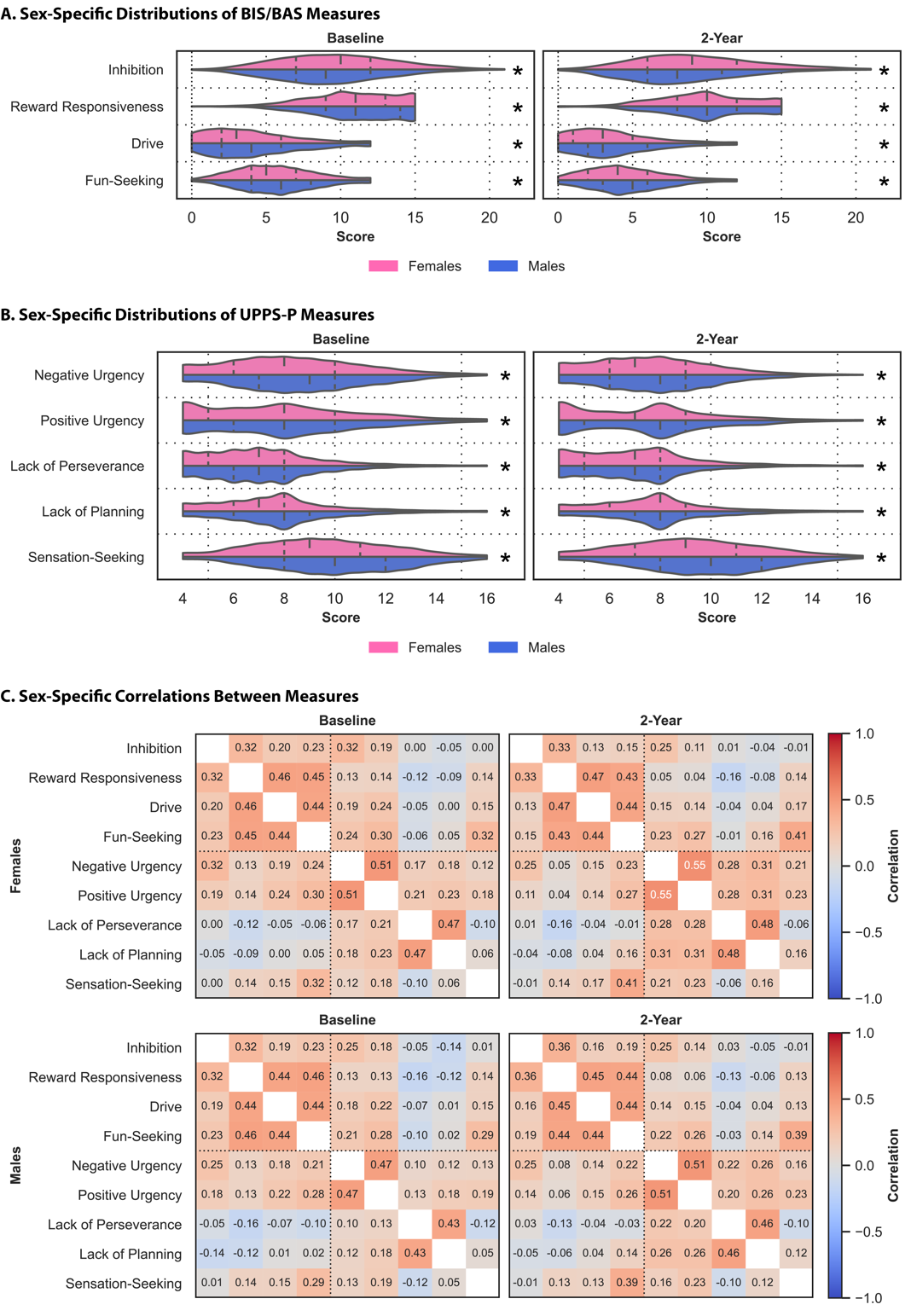

**Figure S6**: **There are sex differences in impulsivity.**
Sex-specific distributions of behavioral measures from the BIS/BAS (A) and UPPS-P (B) scales in females (pink) and males (blue). The shape of the violins indicates the distribution of values, the dashed lines indicate the median, and the dotted lines indicate the interquartile range. Asterisks (*) indicate sex differences. Sex-specific correlations between behavioral measures (C) for females (top) and males (bottom). Baseline (left) and two-year follow-up data are shown. Warmer colors indicate a stronger positive correlation, cooler colors indicate a stronger negative correlation.

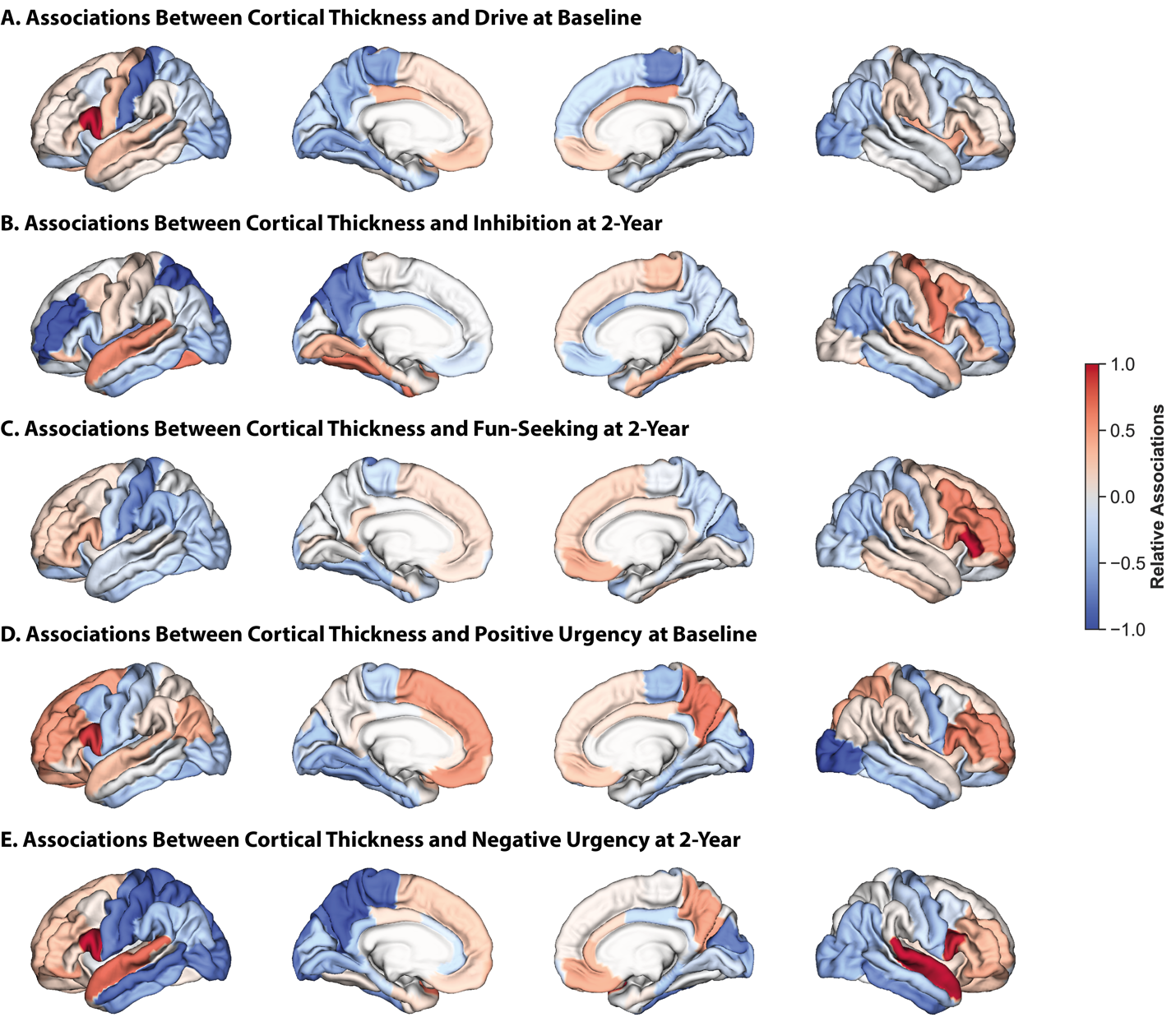

**Figure S7:** **Associations between cortical thickness and reward/punishment sensitivity and impulsivity are dispersed.**
Relative regional associations from models trained on cortical thickness data to predict drive at baseline (A), inhibition at two-year follow-up (B), fun-seeking at two-year follow-up (C), positive urgency at baseline (D), and negative urgency at two-year follow-up (E). Lateral (outer) and medial (inner) surfaces for left (left) and right (right) hemispheres are shown. Warmer colors indicate a stronger positive association, cooler colors indicate a stronger negative association. To facilitate visualization, association values for each set of models were divided by the maximum value for that model.

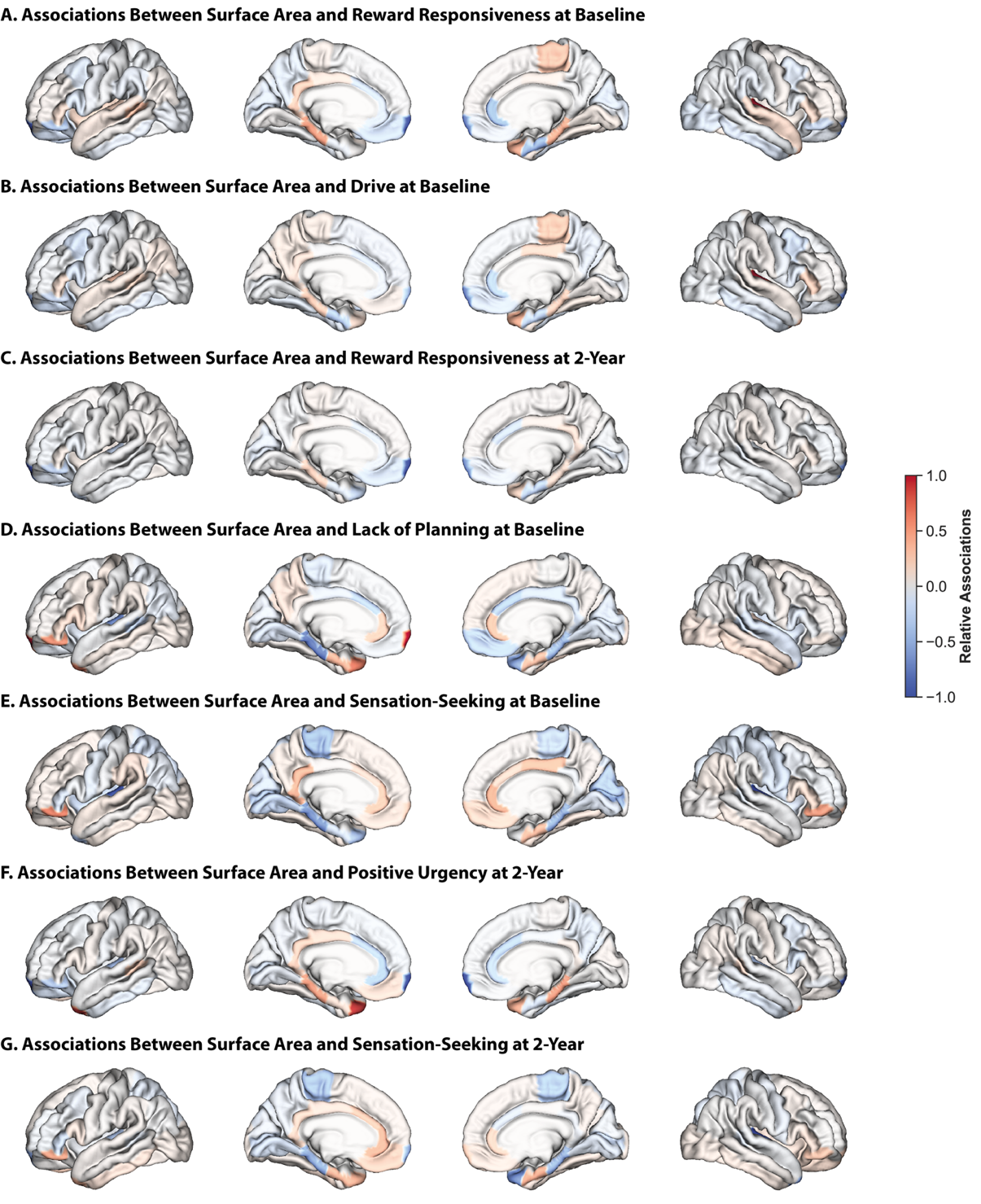

**Figure S8:** **Associations between surface area and reward/punishment sensitivity and impulsivity are localized.**
Relative regional associations from models trained on surface area data to predict reward responsiveness at baseline (A), drive at baseline (B), reward responsive at two-year follow-up (C), lack of planning at baseline (D), sensation-seeking at baseline (E), positive urgency at two-year follow-up (F), and sensation-seeking at two-year follow-up (G). Lateral (outer) and medial (inner) surfaces for left (left) and right (right) hemispheres are shown. Warmer colors indicate a stronger positive association, cooler colors indicate a stronger negative association. To facilitate visualization, association values for each set of models were divided by the maximum value for that model.

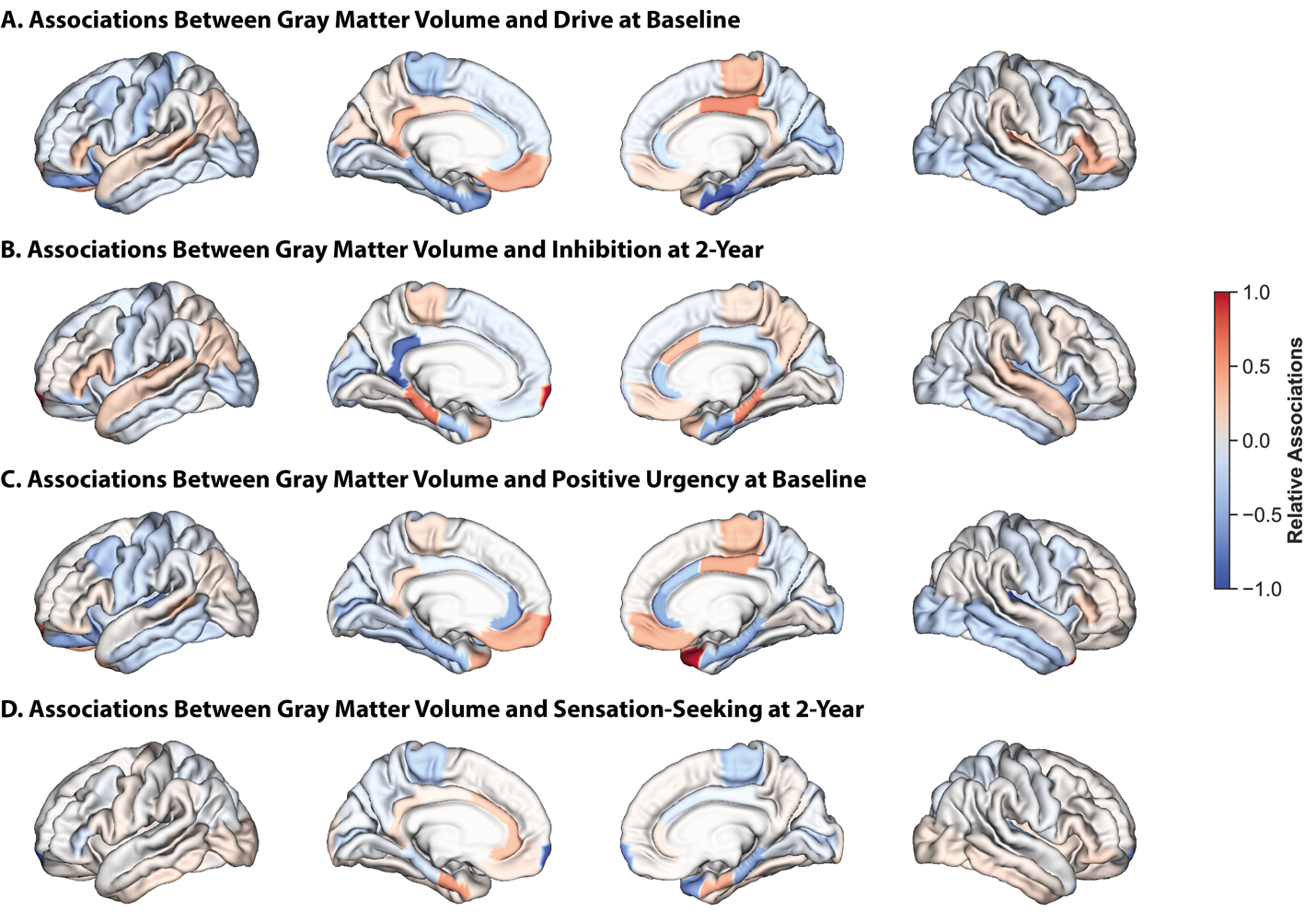

**Figure S9.** **Associations between gray matter volume and impulsivity are localized.**
Relative regional associations from models trained on gray matter volume data to predict drive at baseline (A), inhibition at two-year follow-up (B), positive urgency at baseline (C), and sensation-seeking at two-year follow-up (D). Lateral (outer) and medial (inner) surfaces for left (left) and right (right) hemispheres are shown. Warmer colors indicate a stronger positive association, cooler colors indicate a stronger negative association. To facilitate visualization, association values for each set of models were divided by the maximum value for that model.

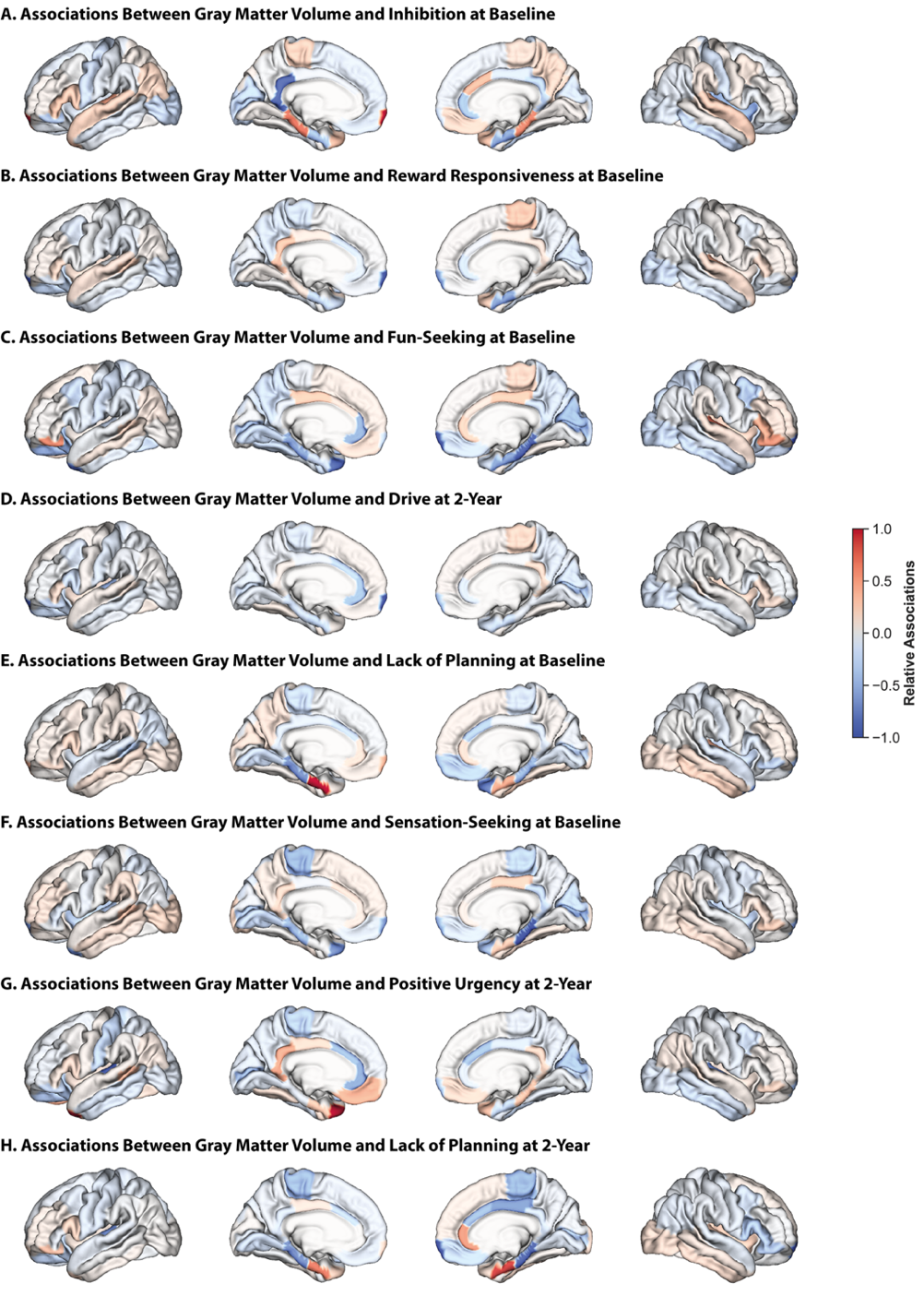

**Figure S10:** **Associations between gray matter volume and reward/punishment sensitivity and impulsivity are localized.**
Relative regional associations from models trained on gray matter volume data to predict inhibition at baseline (A), reward responsiveness at baseline (B), fun-seeking at baseline (C), drive at two-year follow-up (D), lack of planning at baseline (E), sensation-seeking at baseline (F), positive urgency at two-year follow-up (G), and lack of planning at two-year follow-up (H). Lateral (outer) and medial (inner) surfaces for left (left) and right (right) hemispheres are shown. Warmer colors indicate a stronger positive association, cooler colors indicate a stronger negative association. To facilitate visualization, association values for each set of models were divided by the maximum value for that model.

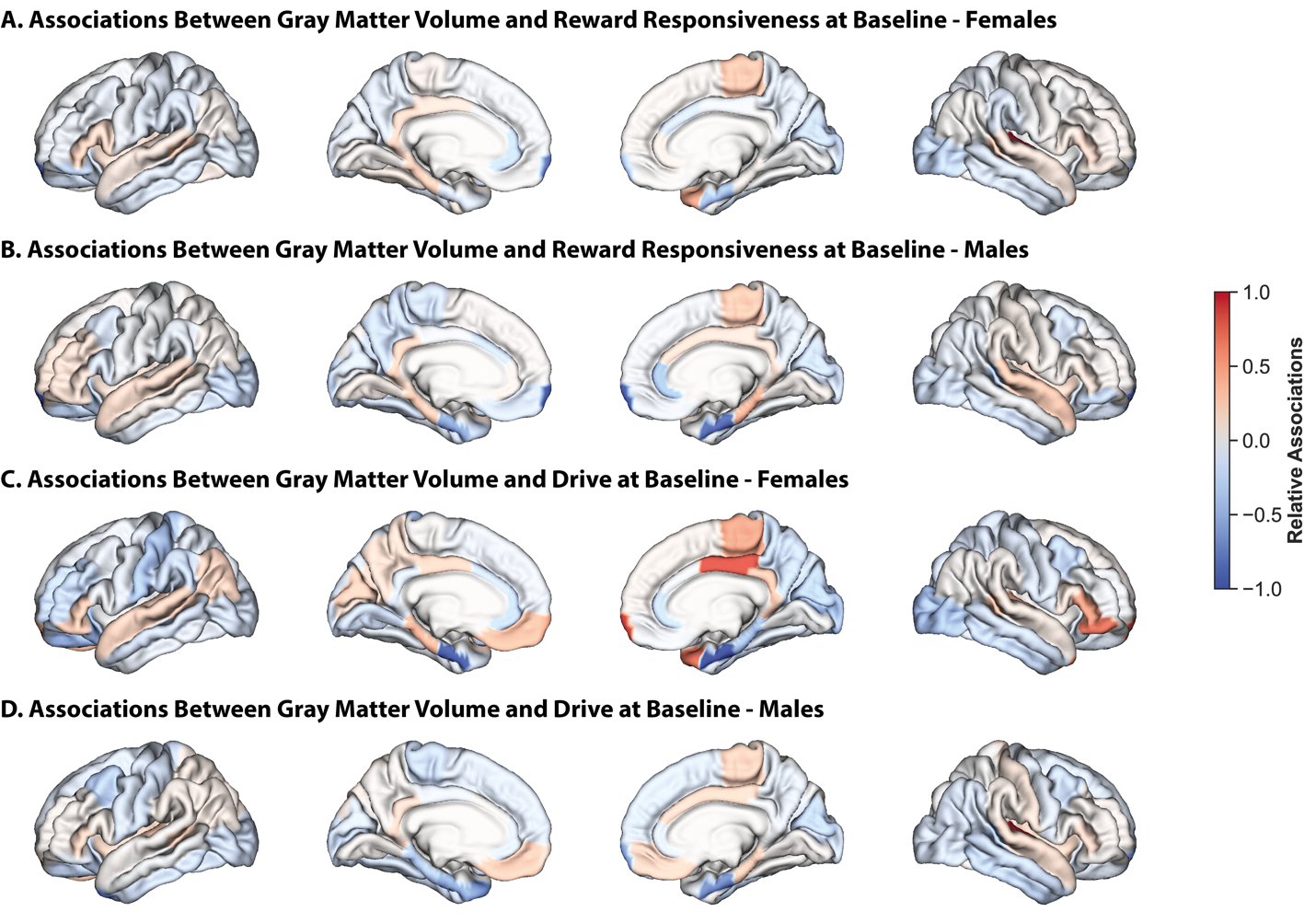

**Figure S11: Associations between gray matter volume and reward responsive are largely shared across the sexes, but those between gray matter volume and drive are considerably distinct.**
Relative regional associations from models trained on gray matter volume data to predict reward responsiveness at baseline in females (A) and males (B), and drive at baseline in females (C) and males (D). Lateral (outer) and medial (inner) surfaces for left (left) and right (right) hemispheres are shown. Warmer colors indicate a stronger positive association, cooler colors indicate a stronger negative association. To facilitate visualization, association values for each set of models were divided by the maximum value for that model.

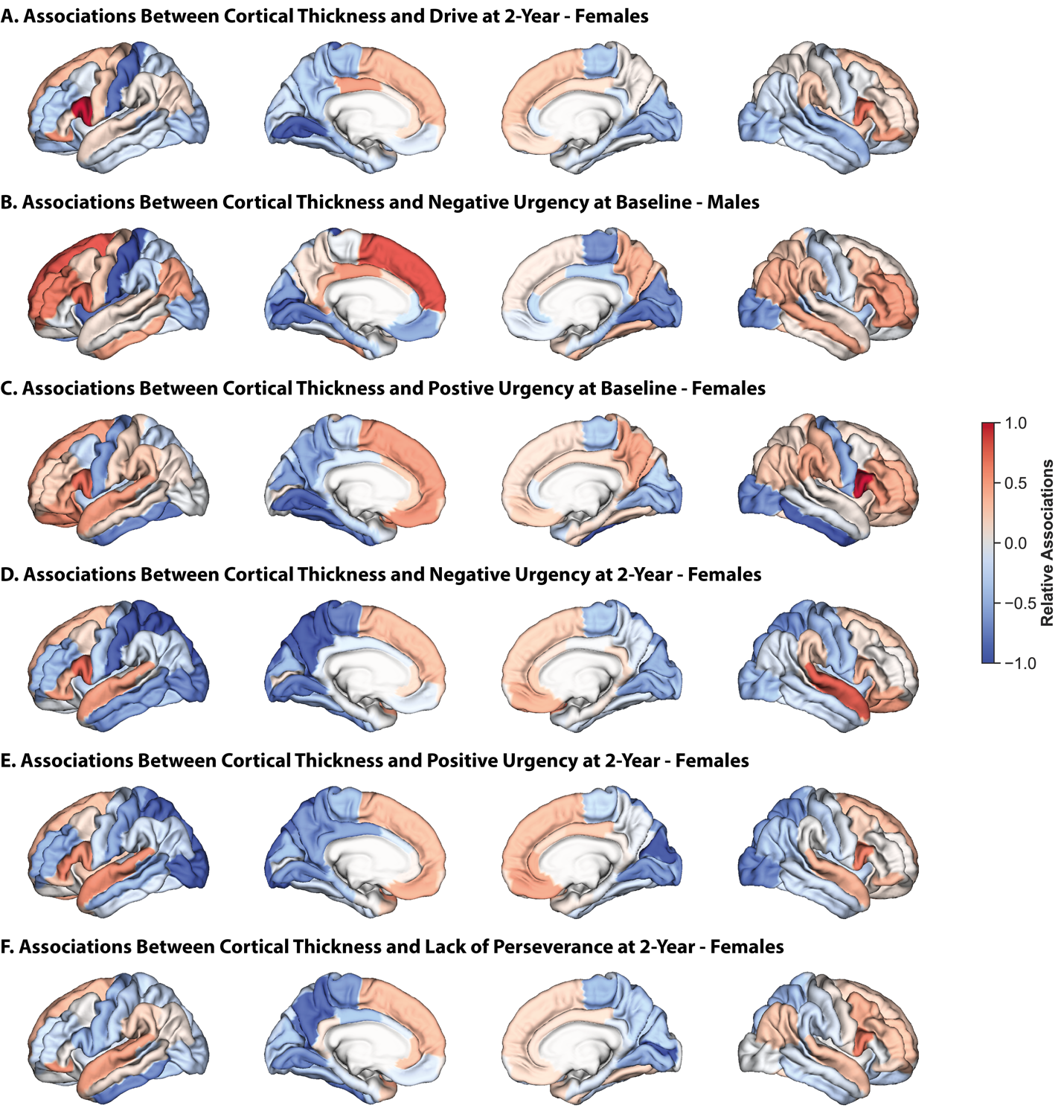

**Figure S12: Results from other sex-specific models based on cortical thickness largely overlap with those from sex-independent models.**
Relative regional associations from models trained on cortical thickness data to predict drive at two-year follow-up in females (A), negative urgency at baseline in males (B), positive urgency at baseline in females (C), negative urgency at two-year follow-up in females (D), positive urgency at two-year follow-up in females (E), lack of perseverance at two-year follow-up in females (F). Lateral (outer) and medial (inner) surfaces for left (left) and right (right) hemispheres are shown. Warmer colors indicate a stronger positive association, cooler colors indicate a stronger negative association. To facilitate visualization, association values for each set of models were divided by the maximum value for that model.

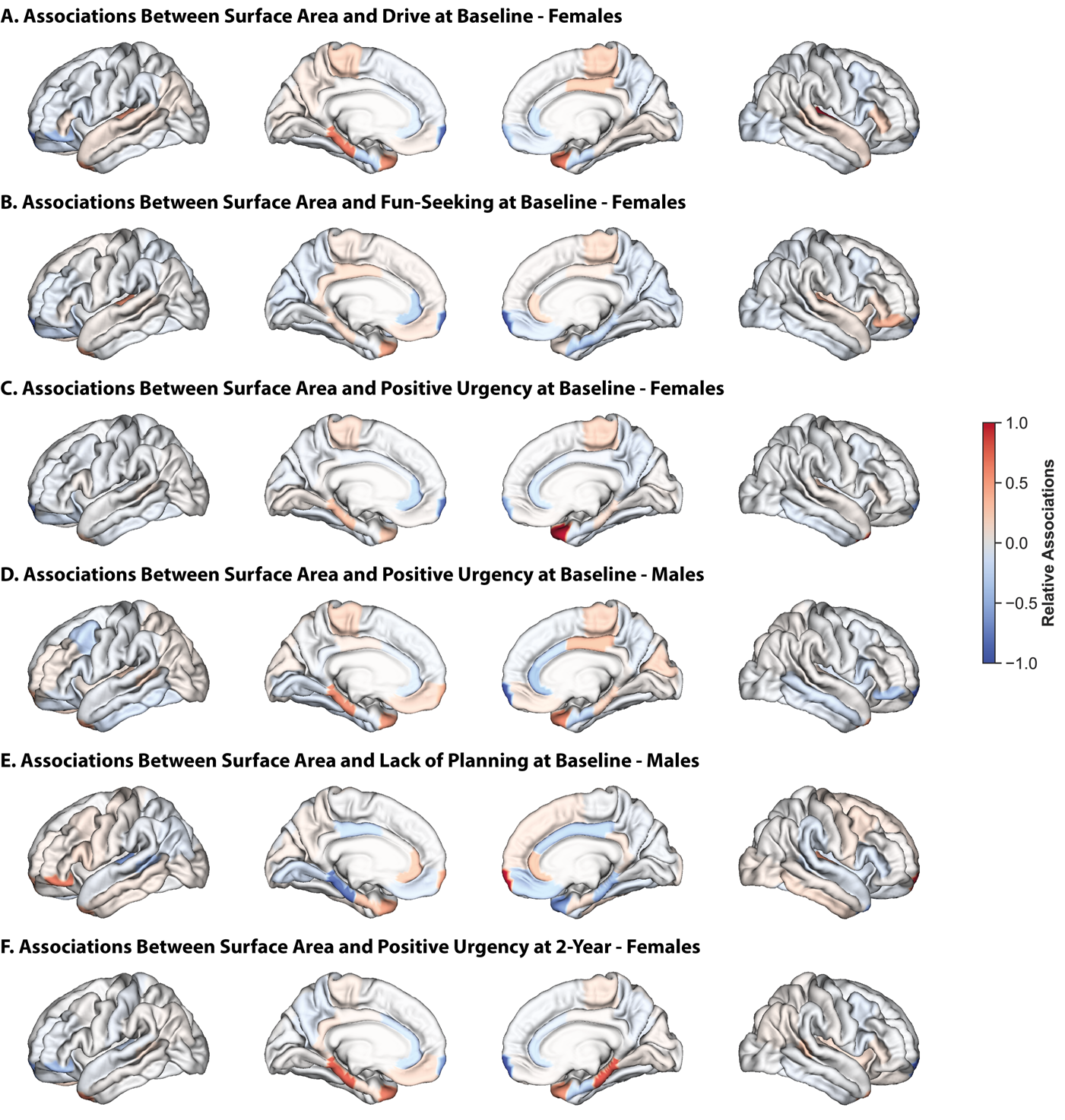

**Figure S13: Results from other sex-specific models based on surface area largely overlap with those from sex-independent models.**
Relative regional associations from models trained on gray matter volume data to predict drive at baseline in females (A), fun-seeking at baseline in females (B), positive urgency at baseline in females (C), positive urgency at baseline in males (D), lack of planning at baseline in males (E), positive urgency at two-year follow-up in females. Lateral (outer) and medial (inner) surfaces for left (left) and right (right) hemispheres are shown. Warmer colors indicate a stronger positive association, cooler colors indicate a stronger negative association. To facilitate visualization, association values for each set of models were divided by the maximum value for that model.

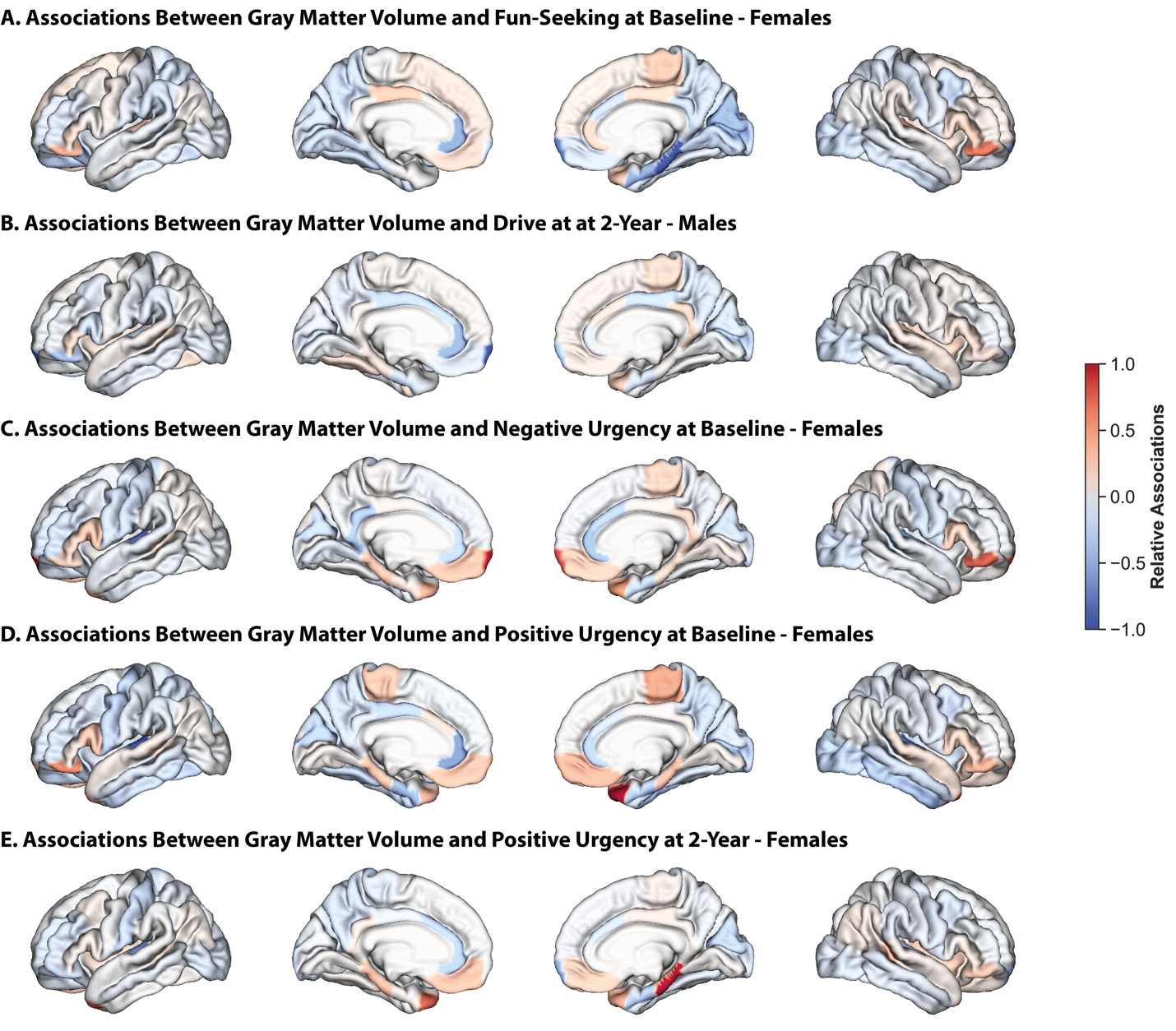

**Figure S14: Results from other sex-specific models based on gray matter volume largely overlap with those from sex-independent models.**
Relative regional associations from models trained on gray matter volume data to predict inhibition at baseline (A), reward responsiveness at baseline (B), fun-seeking at baseline (C), drive at two-year follow-up (D), lack of planning at baseline (E), sensation-seeking at baseline (F), positive urgency at two-year follow-up (G), and lack of planning at two-year follow-up (H). Lateral (outer) and medial (inner) surfaces for left (left) and right (right) hemispheres are shown. Warmer colors indicate a stronger positive association, cooler colors indicate a stronger negative association. To facilitate visualization, association values for each set of models were divided by the maximum value for that model.

|  | **Baseline**  **(n = 9,099)** | **2-Year Follow-Up (n=6,432)** |
| --- | --- | --- |
| **Sex (count, proportion)** | | |
| *Male* | 4759, 52.3% | 3467, 53.9% |
| *Female* | 4337, 47.7% | 2964, 46.1% |
| *Intersex* | 3, 0.0% | 1, 0.0% |
| **Race/Ethnicity (count, proportion)** | | |
| *Asian (4)* | 207, 2.3% | 131, 2.0% |
| *Black (2)* | 1361, 15.0% | 863, 13.4% |
| *Hispanic (3)* | 1944, 21.4% | 1294, 20.1% |
| *Other (5)* | 950, 10.4% | 674, 10.5% |
| *White (1)* | 4636, 51.0% | 3470, 53.9% |
| *No Response* | 1, 0.0% | 0, 0.0% |
| **Socioeconomic Status/Income (count, proportion)** | | |
| *< $,5000* | 324, 3.6% | 185, 2.9% |
| *$5,000 – 11,999* | 339, 3.7% | 207, 3.2% |
| *$12,000 – 15,999* | 220, 2.4% | 130, 2.0% |
| *$16,000 – 24,999* | 407, 4.5% | 281, 4.4% |
| *$25,000 – 34,999* | 521, 5.7% | 349, 5.4% |
| *$35,000 – 49,999* | 701, 7.7% | 530, 8.2% |
| *$50,000 – 74,999* | 1140, 12.5% | 860, 13.4% |
| *$75,000 – 99,999* | 1214, 13.3% | 919, 14.3% |
| *$100,000 – 199,999* | 2503, 27.5% | 1806, 28.1% |
| *> $200,000* | 943, 10.4% | 657, 10.2% |
| *Don’t know* | 388, 4.3% | 264, 4.1% |
| *Refuse to Answer* | 399, 4.4% | 244, 3.8% |

**Table S1: Demographic Information.**
Demographic information (sex, race/ethnicity, socioeconomic status/income) for all participants included at the baseline and 2-year follow-up time points. Reported proportions (%) may not sum to exactly 100% due to rounding.

|  | **Baseline**  **(n = 9,099)** | | | **2-Year Follow-Up (n=6,432)** | | |
| --- | --- | --- | --- | --- | --- | --- |
|  | Cortical Thickness  (r, p_FDR_) | Surface Area  (r, p_FDR_) | Gray Matter Volume  (r, p_FDR_) | Cortical Thickness  (r, p_FDR_) | Surface Area  (r, p_FDR_) | Gray Matter Volume  (r, p_FDR_) |
| **Reward/Punishment Sensitivity** | | | | | | |
| Inhibition | 0.014  (0.328) | 0.020  (0.052) | **0.024**  **(0.023)** | **0.059**  **(0.022)** | **0.037**  **(0.044)** | **0.058**  **(0.010)** |
| Reward Responsiveness | **0.044**  **(<0.001)** | **0.043**  **(<0.001)** | **0.065**  **(0.002)** | 0.005  (0.638) | **0.019**  **(0.044)** | 0.015  (0.129) |
| Drive | **0.093**  **(<0.001)** | **0.064**  **(0.004)** | **0.088**  **(0.004)** | **0.073**  **(0.004)** | 0.045  (0.051) | **0.077**  **(<0.001)** |
| Fun-Seeking | 0.022  (0.284) | **0.035**  **(<0.001)** | **0.043**  **(0.002)** | **0.029**  **(0.028)** | 0.002  (0.271) | 0.008  (0.129) |
| **Impulsivity** | | | | | | |
| Negative Urgency | **0.055**  **(0.020)** | 0.021  (0.059) | 0.028  (0.086) | **0.049**  **(0.008)** | -0.014  (0.483) | 0.017  (0.155) |
| Positive Urgency | **0.078**  **(0.015)** | **0.074**  **(<0.001)** | **0.071**  **(<0.001)** | **0.062**  **(<0.001)** | **0.055**  **(0.040)** | **0.059**  **(<0.001)** |
| Lack of Perseverance | 0.020  (0.166) | -0.016  (0.720) | -0.004  (0.646) | 0.050  (0.082) | 0.028  (0.483) | 0.043  9).102) |
| Lack of Planning | 0.022  (0.050) | **0.061**  **(<0.001)** | **0.062**  **(<0.001)** | 0.016  (0.259) | **0.050**  **(0.040)** | **0.056**  **(0.030)** |
| Sensation-Seeking | 0.026  (0.166) | **0.055**  **(0.003)** | **0.036**  **(0.033)** | 0.032  (0.418) | **0.080**  **(0.025)** | **0.065**  **(0.005)** |

**Table S2: Model Performance for Sex-Independent Models.**
Model performance measures for sex-independent models trained to predict reward/punishment sensitivity and impulsivity based on cortical thickness, surface area, and gray matter volume at baseline and 2-year follow-up. Mean prediction accuracy (r, correlation between observed and predicted values) and FDR-corrected p-values are shown. Models that yielded significant results are denoted by bold text.

|  | **Baseline**  **(n = 9,099)** | | | **2-Year Follow-Up (n=6,432)** | | |
| --- | --- | --- | --- | --- | --- | --- |
|  | Cortical Thickness  (r, p_FDR_) | Surface Area  (r, p_FDR_) | Gray Matter Volume  (r, p_FDR_) | Cortical Thickness  (r, p_FDR_) | Surface Area  (r, p_FDR_) | Gray Matter Volume  (r, p_FDR_) |
| **Reward/Punishment Sensitivity** | | | | | | |
| Inhibition | -0.001  (0.716) | 0.001  (0.271) | 0.003  (0.610) | 0.032  (0.119) | 0.011  (0.630) | -0.005  (0.708) |
| Reward Responsiveness | **0.065**  **(0.002)** | 0.070  (0.051) | **0.052**  **(0.028)** | -0.001  (0.403) | 0.003  (0.630) | 0.008  (0.564) |
| Drive | **0.133**  **(<0.001)** | **0.106**  **(<0.001)** | **0.105**  **(<0.001)** | **0.104**  **(0.028)** | 0.054  (0.63) | 0.080  (0.132) |
| Fun-Seeking | 0.03  (0.56) | 0.031  (0.036) | **0.045**  **(0.006)** | 0.040  (0.352) | 0.017  (0.63) | 0.03  (0.132) |
| **Impulsivity** | | | | | | |
| Negative Urgency | 0.071  (0.092) | 0.040  (0.253) | **0.034**  **(0.040)** | **0.065**  **(0.005)** | -0.001  (0.145) | 0.044  (0.195) |
| Positive Urgency | **0.119**  **(<0.001)** | **0.099**  **(<0.001)** | **0.098 (<0.001)** | **0.093**  **<0.001)** | **0.075**  **(0.005)** | **0.094**  **(0.005)** |
| Lack of Perseverance | -0.005  (0.761) | -0.013  (0.717) | -0.025  (0.744) | **0.051**  **(0.038)** | 0.029  (0.211) | 0.051  (0.225) |
| Lack of Planning | -0.012  (0.546) | 0.002  (0.391) | 0.004 (0.265) | 0.007  (0.622) | 0.022  (0.211) | 0.034 (0.225) |
| Sensation-Seeking | 0.017  (0.055) | 0.027  (0.391) | 0.009 (0.454) | 0.005  (0.622) | 0.047  (0.481) | 0.041  (0.388) |

**Table S3: Model Performance for Female-Specific Models**
Model performance measures for female-specific models trained to predict reward/punishment sensitivity and impulsivity based on cortical thickness, surface area, and gray matter volume at baseline and 2-year follow-up. Mean prediction accuracy (r, correlation between observed and predicted values) and FDR-corrected p-values are shown. Models that yielded significant results are denoted by bold text.

|  | **Baseline**  **(n = 9,099)** | | | **2-Year Follow-Up (n=6,432)** | | |
| --- | --- | --- | --- | --- | --- | --- |
|  | Cortical Thickness  (r, p_FDR_) | Surface Area  (r, p_FDR_) | Gray Matter Volume  (r, p_FDR_) | Cortical Thickness  (r, p_FDR_) | Surface Area  (r, p_FDR_) | Gray Matter Volume  (r, p_FDR_) |
| **Reward/Punishment Sensitivity** | | | | | | |
| Inhibition | 0.005  (0.119) | 0.034  (0.065) | 0.014  (0.574) | 0.005  (0.509) | 0.009  (0.831) | 0.026  (0.675) |
| Reward Responsiveness | **0.046**  **(0.008)** | 0.033  (0.065) | **0.059**  **(0.020)** | 0.015  (0.448) | 0.010  (0.644) | 0.015  (0.050) |
| Drive | **0.060**  **(0.004)** | 0.048  (0.065) | **0.072**  **(<0.001**) | 0.047  (0.448) | 0.020  (0.644) | **0.060**  **(<0.001)** |
| Fun-Seeking | 0.014  (0.148) | 0.042  (0.065) | 0.030  (0.529) | 0.010  (0.509) | -0.049  (0.885) | -0.046  (0.820) |
| **Impulsivity** | | | | | | |
| Negative Urgency | **0.048**  **(0.015)** | 0.020  (0.193) | 0.018  (0.284) | 0.022  (0.558) | -0.050  (0.787) | -0.030  (0.806) |
| Positive Urgency | 0.042  (0.055) | **0.046**  **(0.005)** | 0.045  (0.220) | 0.404  (0.558) | 0.041  (0.228) | 0.025  (0.135) |
| Lack of Perseverance | 0.020  (0.203) | 0.003  (0.426) | 0.013  (0.284) | 0.032  (0.505) | 0.026  (0.110) | 0.033  (0.285) |
| Lack of Planning | -0.004  (0.719) | **0.031**  **(0.008)** | 0.036  (0.284) | 0.007  (0.558) | 0.039  (0.110) | 0.038 (0.285) |
| Sensation-Seeking | 0.008  (0.719) | -0.004  (0.729) | -0.038  (0.540) | 0.037  (0.150) | 0.014  (0.787) | 0.027  (0.418) |

**Table S4: Model Performance for Male-Specific Models**
Model performance measures for male-specific models trained to predict reward/punishment sensitivity and impulsivity based on cortical thickness, surface area, and gray matter volume at baseline and 2-year follow-up. Mean prediction accuracy (r, correlation between observed and predicted values) and FDR-corrected p-values are shown. Models that yielded significant results are denoted by bold text.
